# Scaling dictates the decoder structure

**DOI:** 10.1101/2021.03.04.433820

**Authors:** Jingxiang Shen, Feng Liu, Chao Tang

## Abstract

Despite fluctuations in embryo size within a species, the spatial gene expression pattern and hence the embryonic structure develop *in proportion* with embryo size, known as the scaling phenomenon. For morphogen induced patterning of gene expression, the positional information encoded in the morphogen profile is decoded by the downstream genetic network (the decoder). In this paper, we show that the requirement of scaling sets severe constraints on the geometric structure of the decoder, which in turn enables deduction of mutants’ behavior and extraction of regulation information without going into any molecular details. We demonstrate that the *Drosophila* gap gene system achieves scaling in the way consistent with our theory – the decoder geometry required by scaling correctly accounts for the observed gap gene expression pattern in nearly all maternal morphogen mutants. Furthermore, the regulation logic and the coding/decoding strategy of the gap gene system can also be revealed from the decoder geometry. Our work provides a general theoretical framework on a large class of problems where scaling output is induced by non-scaling input, as well as a unified understanding of scaling, mutants’ behavior and gene regulation for the *Drosophila* gap gene system.

## Introduction

In embryonic development, a cell must know its position in space in order to determine its fate. In the early embryo of *Drosophila* and many other cases, this positional information is encoded in a space-dependent signal like morphogen gradient^1-3^. The decoder in the cell (usually a gene network) reads the local value of the signal to infer the cell’s spatial position and express the appropriate genes, as exemplified in the French-flag model^1,4,5^.

Embryogenesis is under precise genetical regulation in general, but the overall embryo size is not. In long germ-band insects like the *Drosophila*, embryo size equals to the egg size, which is strongly affected by the environmental conditions such as temperature^6^, oxygen level^7^, etc. Therefore, the developmental program must be properly designed to ensure individuals of the same species develop in the same correct proportion regardless of the overall size change. Such scaling property is very common^8-12^ and especially so for *Drosophila*^13-15^.

However, for gene patterning guided by morphogens, scaling does not come by itself. A morphogen gradient is formed via diffusion and degradation. It has a decay length determined by the diffusion constant and the degradation rate, which are usually fixed independent of the embryo size. One solution is to introduce additional regulations to make the morphogen gradient scale by itself, i.e., being able to adjust its decay length according to the embryo size. This kind of strategy is adopted by systems like the *Xenopus* germ-layer specification^16,17^ and the *Drosophila* wing disc^18-20^ and dorsoventral axis patterning^21^. For the *Drosophila* anterior-posterior (A-P) segmentation studied here, much effort has been devoted to measuring scaling property of the key morphogen Bicoid (Bcd)^22-25^, but the results indicate that it has a fixed decay length instead of a scaling one.

Downstream of the maternal morphogens, the gaps are the first set of *Drosophila* A-P segmentation genes, whose expression pattern seems to scale with the embryo length almost perfectly. Evidence of scaling gap gene pattern is reported in WT embryos with natural length fluctuations^22^, as well as in artificially selected fly lines with much larger and smaller embryos^26^. In a recent experiment, scaling of the boundary positions of all four trunk gap genes (*hunchback* (*hb*), *Krüppel* (*Kr*), *knirps* (*kni*) and *giant* (*gt*)) are shown to largely preserve even when embryo length is artificially reduced by 30%^25^. Although those results are far from perfect – the measured scaling error is around 3% embryo length, larger than the baseline noise in embryos of identical size^27,28^, we believe that this 3% accuracy is enough to confirm the presence of scaling mechanism in the gap gene system, since the “null hypothesis” that gap gene simply follow the Bcd thresholds should give a scaling error of at least 10% (Fig. S7f, lower right panel). Therefore, although some subsequent refinements may still be possible, much of the scaling job is done at the gap gene stage by reading the non-scaling maternal morphogens.

There are many proposed mechanisms attempting to solve the gap genes scaling problem, e.g., amplitude correction or “partial scaling” of the morphogen Bcd gradient^24,29,30^, decoding unsteady Bcd gradient^31,32^, the dynamical “canalization” of the gap gene network^33^, and positional error correction based on diffusion of the gap gene products^34^. These studies provided much data and insights. Nonetheless, a comprehensive understanding that accounts for scaling throughout the entire embryo is still lacking, as well as predictions that can be validated systematically and experimentally.

Another valuable early idea is the bi-gradient model^1,35,36^ – if a second posterior gradient is present in addition to the anterior Bcd gradient, a cell can in principle have enough information to “compute” its relative position in the embryo. For historical reasons, the bi-gradient model is not well-developed (see SI-12) and remains largely a theoretical possibility for the gap gene case, though this idea that integrative decoding of two opposing gradients has been successfully applied and extended in a later work to vertebrate neural tube development^37^.

The bi-gradient model represents the idea of “local decoder”, i.e., the cell fate at each spatial position is a function of morphogen levels at this spatial point *alone*. The developmental gene regulation network is effectively a decoder, working in a spatially decoupled manner, mapping a combination of local morphogen concentrations to a local cell fate. This decoding idea has been widely adopted in analyzing optimal extraction of positional information from single or multiple noisy morphogen gradients^37-41^. Such “local decoding” framework is the minimum but important starting point for theoretical analyses including ours.

Measuring the morphogen gradients and wild-type gap gene patterns to such a precision that can directly distinguish between the above-mentioned theoretical models is extremely challenging, which makes the gap gene scaling problem remains to be settled. In this paper we approach this problem in a different way. By starting from the assumption that scaling can be achieved within the local decoder framework, we show that the necessary and sufficient condition for a local decoder to generate a scaling gene expression pattern across the embryo contains rich information on the geometric structure of the decoder, i.e., the decoder structure is dictated purely from its function (scaling). Then, the bulk of the existing experiments on morphogen mutants can be employed in the study of scaling – in our framework it is just the same decoder applied to altered inputs. The measured mutants’ patterns are in excellent agreement with the prediction by the scaling local decoder, which strongly support our stating assumption that scaling of the *Drosophila* gap gene system should originate from an integrative decoding of the multiple non-scaling input gradients.

## Results

### The Geometric Structure of the Decoder is Determined by the Scaling Requirement

We first illustrate our basic idea using a bi-gradient model^1,35,36^, and demonstrate how the decoder structure can be dictated directly from scaling requirement. In this simplified model, there are two morphogens, M_1_ and M_2_, having anterior and posterior exponentially shaped gradients with a fixed length constant *λ* (Fig. 1a, length of a standard-sized embryo is used as the length unit).

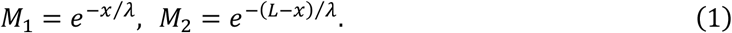

**Fig. 1.**
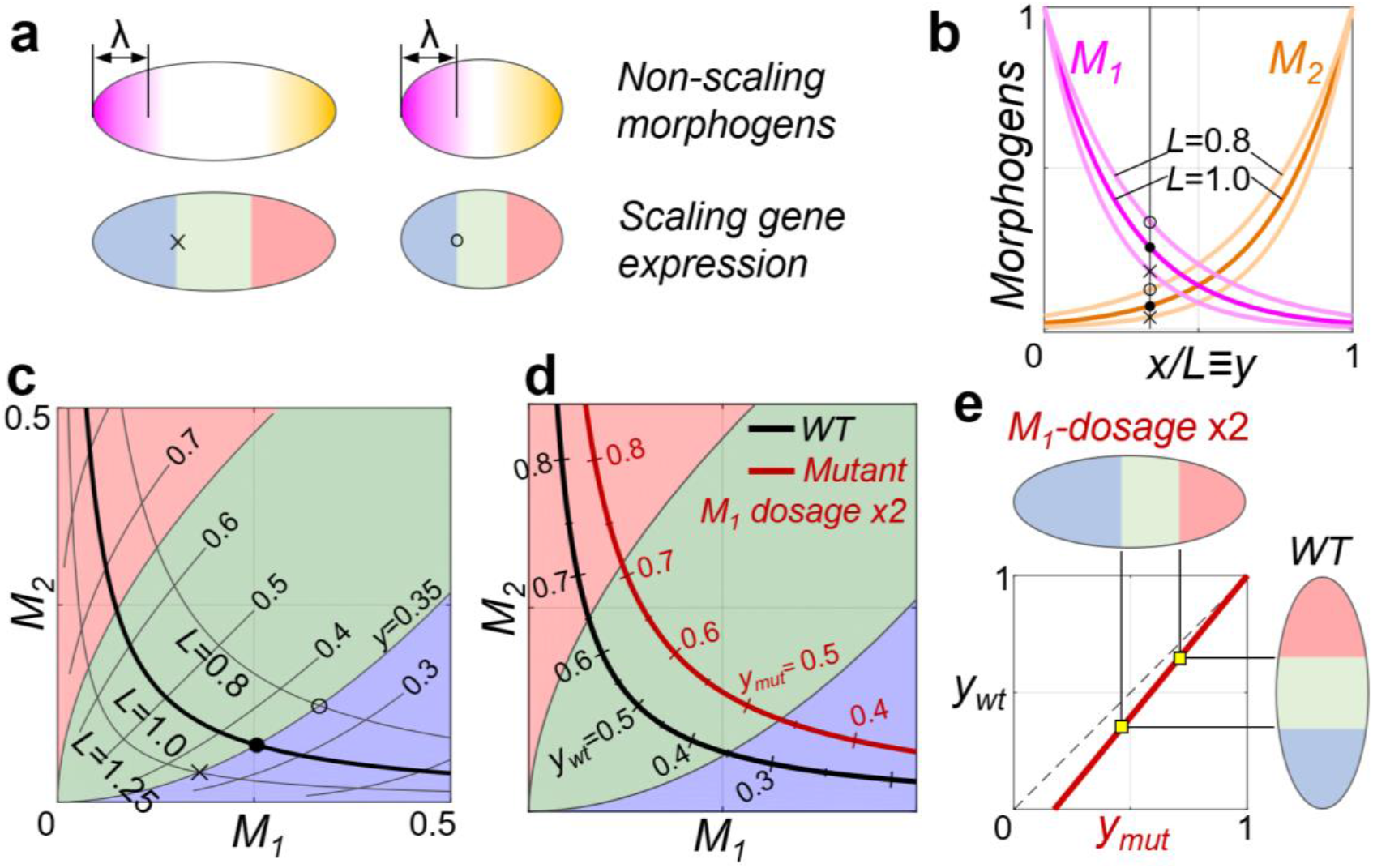
Generating scaling output by reading the local values of two non-scaling gradients. (**a**) A schematic sketch with two embryos of different sizes. The two morphogen gradients are shown in the upper panel and the desired scaling output patterns are shown below. (**b**) The same relative position *y*≡*x/L*=0.35 has higher level of both morphogens in a smaller embryo (marked by “O”) than a larger one (“X”). (**c**) In the M_1_-M_2_ space, the standard-sized WT embryo is represented by the black curve. As *L* varies, the *y*=0.35 point traces out a line. (**d**) Maternal morphogen profiles in a mutant are also represented by a curve in the M_1_-M_2_ space. The red curve stands for an *L*=1 embryo where the M_1_ dosage is doubled. The corresponding cell fates along this red curve, hence along the A-P axis of this mutant, can be directly read out. (**e**) The predicted “fate map” of the mutant in (**d**). All gene expression boundaries, if their positions in WT were plotted against their shifted positions in the mutant, should lie on the fate map.

Define the relative coordinate *y*≡*x/L*, so that the larger (*L*>1) and the smaller (*L*<1) embryos can all be placed together with the standard-sized embryos (*L*=1) for comparison. Scale-invariance of an expression pattern *F*(*x,L*) means that it depends only on the ratio of *x* and *L*: *F*(*x,L*)=*f*(*x*/*L*)=*f*(*y*). Obviously, the morphogen gradients with fixed length constant (Eq. 1) are not scale invariant with respect to the change of the length scale *L* (Fig. 1b),

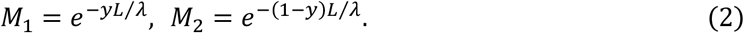

Consider for example the position *y*=0.35 as the boundary separating two different cell fates in the embryo. In a smaller (larger) embryo, as the absolute distance to both termini are shorter (longer) at this position, local levels of both morphogens are higher (lower). When *L* varies, the *y*=0.35 point traces out a line on the M_1_-M_2_ plane (Fig. 1c). Therefore, for a local decoder to give perfect scaling outputs, this *y*-constant line must be followed by the decision boundary of the decoder, which maps a (M_1_, M_2_) pair to a gene expression state (“fate”). In other words, the requirement imposed by scaling is enough to determine the effective input-output relation (coloring scheme) of the decoder in this double-gradient case, no matter how the decoder is implemented biochemically. The cell fate can also be represented by its position in wild-type (WT), denoted as 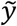 In this case:

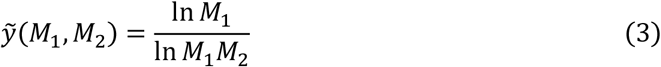

which is obtained by eliminating *L* from Eq. 2.

Once determined by its scale-invariant performance in WT embryos, the same decoder can be applied to mutant embryos where the maternal morphogen profiles are perturbed but the decoding machines are intact. In the case considered here, all *y* points belonging to a WT embryo of size *L* correspond to the hyperbolic curve *M*_1_ · *M*_2_ = *e*^−*L*/*λ*^. On the other hand, consider a standard-sized (*L*=1) mutant embryo where the M_1_ copy number is doubled:

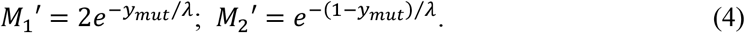

This mutant is also represented by a hyperbolic curve *M*_1_′ · *M*_2_′ = 2*e*^−1/*λ*^ (Fig. 1d). Note that this curve is exactly the one corresponding to a WT embryo of size *L* = 1 − *λ* ln 2, though points of the same cell fate locate at different *y*’s in WT and the mutant. For example, the blue-green boundary at *y*=0.35 in WT is shifted to 0.46 in the mutant.

In general, for any *y*_*mut*_ in this mutant embryo, there always exists a corresponding position *y*_*WT*_ in WT embryo with the same morphogen values, hence the same cell fate 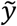. In this case, substituting the mutant’s morphogen profile (Eq. 4) into the decoder (Eq. 3) yields such a mapping:

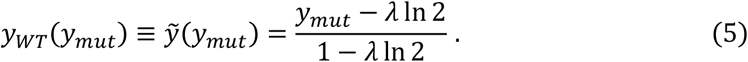

We call this mapping 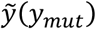 the “fate-map” of this mutant. This fate map can be tested experimentally by plotting the positions *y*_*WT*_ against *y*_*mut*_ for the measured expression domain boundaries (Fig. 1e). If the measured gene profile in mutants match the scaling predictions, then it would be a strong support that the underlying gene network decodes the multiple morphogens in such a specific way to enable scaling. In the following sections we demonstrate that the *Drosophila* gap gene system is indeed a decoder of this kind – scaling and mutants’ behavior are just different aspects of the same underlying decoder geometry.

### Construction of a Phenomenological Decoder for *Drosophila*

Following the procedure outlined above, we next construct a decoder for the *Drosophila* gap gene system, using all the three maternal morphogen gradients Bcd, Nanos (Nos) and Torso (Tor). These three are also the only upstream-most gradients^42^, since *bcd*^*-*^*nos*^*-*^*tor*^*-*^ triple mutant shows spatially uniform gap gene expression^40,43^. Other gradients (like Caudal) is known to be derived from these three primary morphogens, thus should not be considered explicitly.

Experimental measurements suggest that Bcd has a fixed length constant^13,22,24,25^ and an overall amplitude positively correlated with embryo length^24,29,30^. So, it is modeled as:

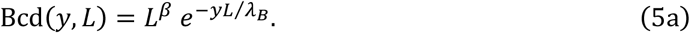

It is evident from Eq. 5a that there is a special point *y* = *βλ*_*B*_ where the Bcd level is *L*-independent owning to the amplitude correction effect. Some previous studies claimed that this effect makes Bcd a “partial scaling” gradient, and may be the main reason for gap gene scaling^24,26^. Although we do not agree with this explanation in general, as it cannot account for scaling throughout the entire embryo especially in the head and abdomen regions far away from the *y* = *βλ*_*B*_ point^14,15,25,26^, this amplitude factor has been verified experimentally, thus should be considered here if we wish our decoder correctly describe the real situation in *Drosophila*.

The *Drosophila* posterior gradient Nos is also formed through localized synthesis and diffusion and has an exponential profile fixed to the posterior: 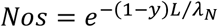. It is well known that Nos functions solely through repressing the maternal component of the gap gene product Hb (mHb) in the posterior half of the embryo^44-46^. Therefore, the “immediate” posterior morphogen should be mHb. If the inhibition of mHb by Nos is modeled by an inhibitory Hill function, then the mHb gradient takes a sigmoidal shape:

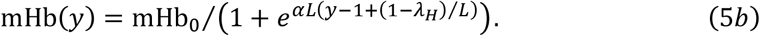

The source of Tor activity locates at both termini of the embryo and extends spatially through diffusion^47,48^. By minimal assumptions, we take a fixed length constant for its profile.

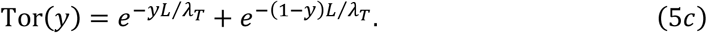

All parameters above are fitted from published experimental data^24,25,44-46,49,50^ (see SI-1 for details). Taken together, in parallel with Eq. 2, Eq. 5 gives a complete description on the *Drosophila* morphogens responsible for A-P patterning, as shown graphically in Fig. 2a.

**Fig. 2.**
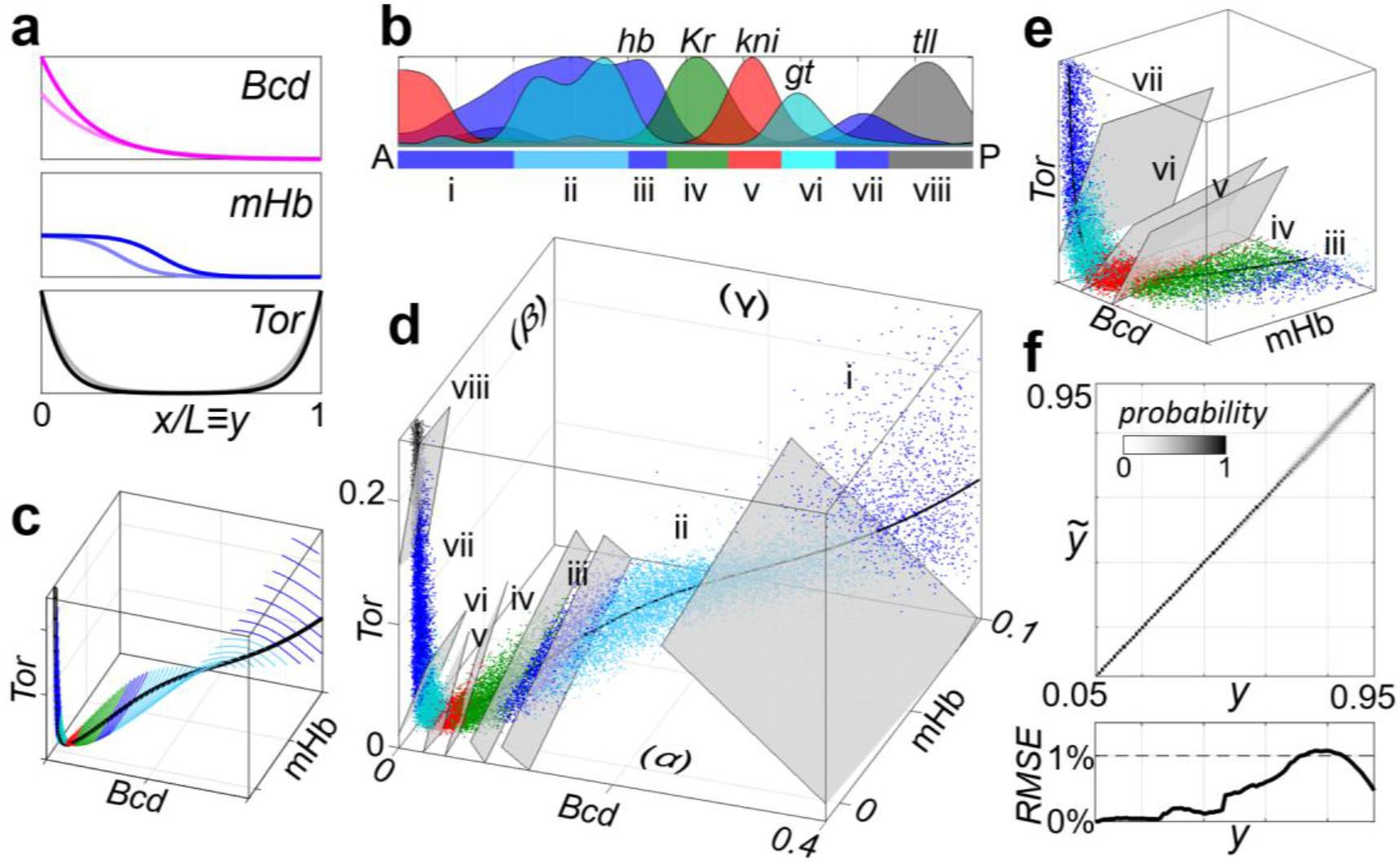
The phenomenological scaling decoder for *Drosophila* A-P patterning. (**a**) The three non-scaling maternal gradients in the relative coordinate *y*≡*x/L* (Eq. 5). The morphogen profiles for a standard size WT embryo (*L*=1, darker lines) and a smaller one (*L*=0.8, lighter lines) are shown. (**b**) About 100 different cell fates along the A-P axis are grouped into 8 domains according to the gap gene expression. The normalized gap gene profiles are adopted from Ref.^27^ and a Gaussian smoothing is applied. (**c**) In the (Bcd, mHb, Tor) space, the standard-sized WT embryo is represented by the black curve along which *y* varies from 0 to 1. When *L* changes, each point on this curve traces out a line representing the morphogen values at this *y* position in WT embryos of different sizes. Only the region corresponding to *L*=0.8∼1.2 are plotted here. The *y*-constant lines shown there have spacing *Δy*=0.01 along the A-P axis. (**d**) A Poisson noise is added to the morphogen levels and the embryo length is sampled from a normal distribution, turning the 2-d WT manifold in (C) to a point cloud. The optimal decision boundaries can be well approximated by a set of linear planes. (**e**) A magnification of (**d**). The iv-v, v-vi and vi-vii boundary planes are shown. (**f**) Positions decoded from the morphogen values 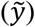 *vs*. the ground truth (*y*) for the WT point cloud in (**d**). Since gap gene expression is affected by the Dorsal-Ventral system when being very close to embryo termini, we only discuss *y* between 0.05 and 0.95 hereafter. The root-mean-square-error is hardly larger than 1%.

Downstream of the maternal morphogens, the gap genes (*hb, Kr, kni* and *gt*) are the first to display a scaling pattern^22,26^. The *Drosophila* embryo has around 100 rows of cells along the A-P axis before gastrulation. Nearly any two of them can be distinguished by their gap gene expression^28^, so there are effectively around 100 different cell fates along the A-P axis and the fate map 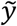 should be discussed at the resolution of 0.01. To visualize these cell fates by colors, we group them into 8 classes according to the dominantly expressing gap gene (Fig. 2b).

A standard-sized (*L*=1) WT embryo is represented by a 1-d curve in the space of (Bcd, mHb, Tor). When *L* varies, this curve sweeps out a 2-d WT manifold. Following our arguments in Fig. 1, the ideal output of a scaling decoder on this WT manifold can be immediately determined – lines of constant *y* values should have the same cell fate (Fig. 2c).

In reality, *Drosophila melanogaster* embryo length varies approximately between 450 and 570 μm (∼±10%) across different fly stocks ^14^, while scaling for the peaks and boundaries of the gap gene expression is near perfect at least to the resolution of currently available measurements^13,15^. To have a more realistic picture, we hereafter sample *L* from a normal distribution *L* ∼ 𝒩(*μ* = 1, *σ* = 0.1) to simulate the effects of length fluctuation across *Drosophila melanogaster* stocks. Another important aspect is noise in morphogen profiles. When morphogen level approaches zero it should have huge relative noise thus cannot carry useful information. Also, there are embryo-to-embryo variability in morphogen amplitude. We introduce explicitly Poisson noise terms to the morphogen values to reflect that the gradients themselves are noisy (See Methods and Supplemental text S2 for details). Amplitude of such noise is selected to reflect the experimentally observed variations. For example, Bcd is the most extensively studied gradient, whose positional error at each spatial position has been carefully measured^51^. Our Poisson noise term is then selected to introduce a positional error of the same order (Fig. S2). As a result, the “colored lines” (i.e., ideal decoder output) in Fig. 2c now transform into more realistic colored point-clouds in Fig. 2d.

The gap gene network (the decoder) should be able to classify all the points in Fig. 2d into classes (real cell fate) that matches their colors. In a sense, this is an “optimal decoding” problem in the presence of noise^28,37,38,40^. Here, the dominant source of noise is the embryo length variation and the main goal is to preserve scaling.

Besides just obtaining a decoder that performs such classification task (like a Bayesian one discussed in Supplemental text S5), we are more interested in figuring out the shape of those decision boundaries explicitly. Complexity and nonlinearity of the decision boundaries are directly related to whether it can be satisfactorily approximated by a gene regulation network, as will be discussed later. Here comes an important observation, that for the *Drosophila* case the boundaries between regions of different desired outputs (colors) are effectively linear – any local decoder that achieves scaling must effectively behave like a set of *linear* classifiers, at least within the WT point clouds. Therefore, we fit the phenomenological decision boundaries with planes (see Methods and Supplemental text S3 for details). Points of different colors seem to be separated satisfactorily in this way (Fig. 2d, e, and Fig. S5d).

The three maternal gradients can not only determine boundaries of gap gene expression domains, but also ∼100 distinct A-P cell fates as mentioned above. Thus, we can construct a (approximately) continuous version of the decoding function using more linear classifiers of this kind. Along the A-P axis, we fit 100 such classification planes at 100 equally spaced *y* positions (Methods and Supplemental text S3). The linear classifiers work sufficiently well with the decoding task, i.e., possible nonlinearities in the ideal classification boundaries can indeed be safely ignored. Fig. 2f shows the decoding result of the WT ensemble of Fig. 2d using the linear classifiers. Despite the presence of Poisson noise, positions 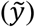 decoded by reading the maternal morphogen values are always close to the ground truth positions (*y*). The root-mean-square-error 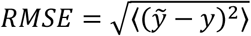 is hardly larger than 1% (reminding that the morphogen noise level here are already comparable to the measured values), even smaller than the *RMSE* of a Bayesian decoder which allows arbitrary decision boundary geometry (Fig. S5a-c).

### The Decoder Quantitatively Predicts Phenotypes of Morphogen Mutants

We next demonstrate that such a decoder, whose overall geometry is solely determined by scaling, has a remarkable power to predict nearly all phenotypes of maternal morphogen mutants in *Drosophila*, demonstrating it to be a satisfactory description of the actual gap gene system. Note that the values of (Bcd, mHb, Tor) in mutants may lie outside of the WT region (color point clouds in Fig. 2d). In this case, we linearly extrapolate the classification planes (see Supplemental text S4 for justification of the extrapolation).

Fig. 3a-c show the intersections of the linearly extrapolated classification planes with the bottom /left /back faces of the cube in Fig. 2d (marked by α /β /γ, on which Tor=0 /Bcd=0 /mHb=mHb_0_), with the same color scheme as in Fig. 2. Maternal morphogen null mutants lie on these faces. For example, in the *nos*^*-*^ mutant mHb is equal to mHb_0_ throughout the entire embryo, corresponding to projecting the WT curve onto the *γ* plane. It is clear in Fig. 3c that along this projected *nos*^*-*^ curve, domain iv is followed immediately by domain vii, indicating the loss of abdominal *kni* (v) and *gt* (vi) domains. This is exactly the case observed in experiments^40,52,53^.

**Fig. 3.**
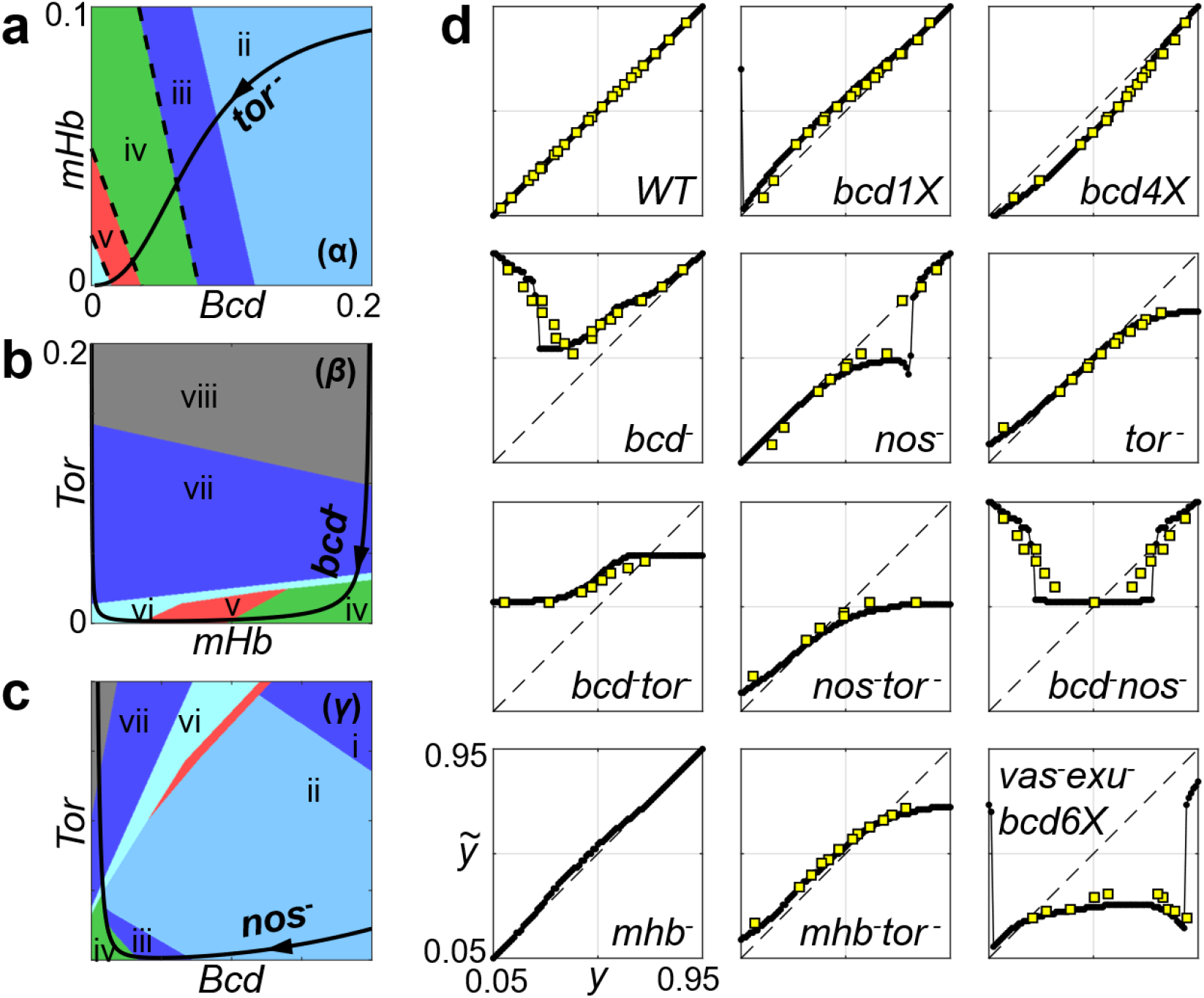
Quantitative predictions on mutant fate maps by the phenomenological decoder. (**a**-**c**) The decoding results on the bottom/left/back faces (marked by α/β/γ) of the cube in Fig. 2d. Solid black curves are projections of the *L*=1 WT curve onto these planes, which also represents the standard-sized mutant embryos *tor*^*-*^/*bcd*^-^/*nos*^-^, respectively. The predicted gap gene expression in these mutants can be read out along these lines (arrowheads on them are pointing from head to tail). (**d**) Fate map 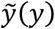 predicted for WT and another 11 different mutants (black lines) and the comparison with experimental measurements (yellow squares, cited from Refs.^27,40,50,54,55^). See Supplemental text S6 for discussion on the panels of *mhb*^*-*^ and *vas*^*-*^*exu*^*-*^*bcd6X*.

As another example, consider the *bcd*^*-*^*tor*^*-*^. mHb is now the only morphogen gradient, decreasing from its maximum value to zero from the anterior to posterior pole. It is obvious in Fig. 3a, b that points on the mHb axis fall into the iv, v and vi domains successively, corresponding to three gap gene domains *K*r, *kni* and *gt* appearing successively in this mutant embryo. As the Bcd=Tor=mHb=0 point is classified into domain vi, the *gt* domain should extend all the way to the posterior pole. This is also the pattern observed experimentally^40,43^.

More than predictions on the presence or absence of certain gap gene domains, a nearly continuous valued fate map 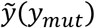 can be constructed by incorporating all the 100 classification planes. That is, the 100 decision planes divide the morphogen space into 101 slices, corresponding to cell fates 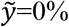 to 100% (See Methods for a detailed description of the procedure). In Fig. 3d, the predicted fate map for WT and 11 maternal morphogen mutants are shown as black curves. To see if these predictions from scaling match experiments, we identify the peak and boundary positions for all gap gene domains (Table S1) from the published quantitative measurements of gap gene profiles in WT and various mutants^27,40,50,54,55^. For each of them, its position in mutant is plotted against its WT positions (yellow squares in Fig. 3d). Remarkably, predictions and experiments agree quantitatively in all cases.

Predicted fate map can also be converted to a predicted gap gene pattern. For *y*_mut_ in a mutant embryo, the gap gene expression level can be approximated by the composite function 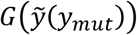, where *G* stands for the WT gap gene expression pattern, and 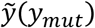 is the fate map. Fig. 4a, b show two examples (solid lines). Measured profiles^40^ are shown as dotted lines in lighter colors for comparison.

**Fig. 4.**
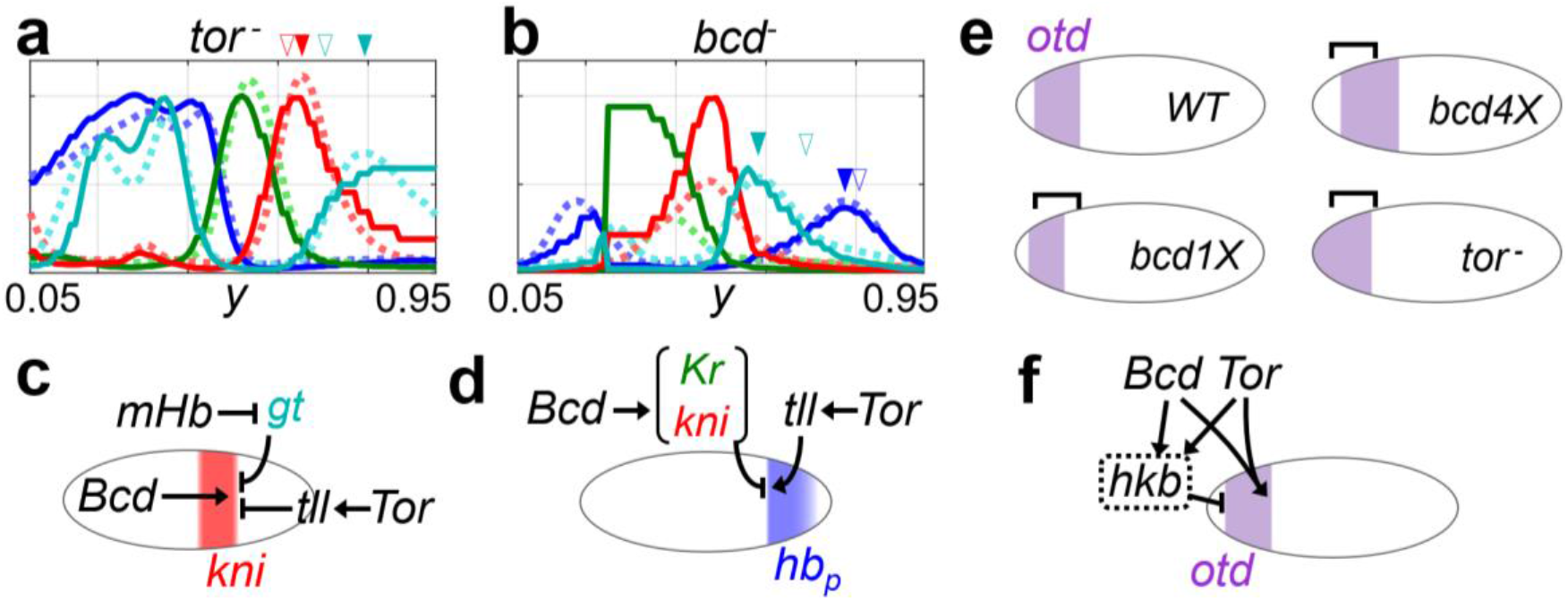
The decoder reveals information about gene regulation logic. (**a**-**b**) The predicted gap gene patterns (solid lines) in *tor*^*-*^ and *bcd*^*-*^ are compared to that measured in Ref.^40^ (dashed lines). The peak and boundary positions are correctly predicted. Peak position of the posterior *kni, hb* and *gt* domains are marked by filled arrowhead, their WT positions by the empty arrowheads. (**c**) “Redundant” regulation on the posterior boundary of *kni* domain as required by scaling. (**d**) According to the decoder, the vi-vii boundary 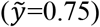 is set by the balance between the opposing gradients Bcd and Tor. Biochemically, this is realized by an indirect activation of Tor through *tll*, together with an indirect inhibition from Bcd, meditated by *Kr* and *kni*. (**e**) The predicted expression patterns of the head gap gene *otd*, derived from the decoder. Structure of the decoder requires both Bcd and Tor to effectively activate *otd* for positioning its posterior boundary, while at the same time, both inhibit *otd* for defining its anterior boundary. This explains the *otd* regulations proposed by experimentalists decades ago (**f**).

### Decoder Geometry Constrains Structure of Underlying Gene Regulation Network

Further analysis reveals interesting connections between the decoder structure and the underlying gene regulation. That is, the *Drosophila* gap gene network is structured in a way that enables scaling.

Consider the *tor*^*-*^ mutant as an example. As dictated by scaling, the decision boundaries *Kr*-*kni, kni*-*gt* and *gt*-*hb* should be inclined in the 3-d morphogen space, not perpendicular to the Bcd-mHb plane (Fig. 2e). This means that Tor should participate in positioning these boundaries to allow for scaling. Geometrically, when extrapolated to the Tor=0 plane following these inclined classification planes, a cell fate in WT should appear at a more posterior position than if being orthogonally projected downward. Thereby in the *tor*^*-*^ mutant, besides the fact that the posterior *hb* domain disappears as a result of lacking the activation from *tailless*^56^, the rest of the abdominal domains should also shift posteriorly. This prediction is fully consistent with experiments (Fig. 4a, where the filled and empty triangles mark the measured peak positions in *tor*^*-*^ and WT, respectively).

This example may help us to understand the ubiquitous “redundant” regulations in the gap gene network. According to the simplest interpretation, mHb gradient defines the anterior boundary of abdominal *gt* (vi) domain through inhibition; Gt inhibits *kni*, thereby setting the posterior boundary of *kni* (v) domain^43^. There is no “need”, in principle, for Tor to be involved. But the observed shift of *kni* boundary in *tor*^*-*^ clearly shows that in reality Tor contributes (probably through *tll*) to the repression on *kni*^53,57,58^, setting its posterior boundary together with mHb as well as Bcd^59^. We propose that such seemingly redundant regulations in fact tune the slope of *kni*-*gt* classification plane, so that it could align with the angle required by scaling (Fig. 4c). Scaling seems to be one important “goal” of such redundancy in the gap gene network.

Another example deals with the *bcd*^*-*^ mutant (Fig. 4b). Bcd is well known to function in the anterior part. However, the region where Bcd plays a role seems to be much wider than naively expected – *bcd*^*-*^ mutant affects even domain vi and vii near the posterior pole as shown by experiments. This aspect of Bcd should also contribute to tuning the decision boundary orientations to allow for scaling, as it has clearly been captured by our scaling-based decoder. To be more precise, the decision plane representing the *gt-hb* (vi-vii) boundary at 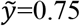 is:

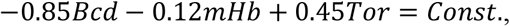

indicating that Tor should be effectively an activator for *hb* here while Bcd should play a repressive role (Fig. 4d). This speculation is consistent with the existing biological knowledge: Tor is known to activate the posterior *hb* domain through *tailless*^57^. The effective repression by Bcd is probably meditated by *Kr* and *kni*, upon whose mutation the posterior *hb* domain expands anteriorly^60^.

A third example of this kind regards the head gap gene *orthodenticle* (*otd*), which is expressed between 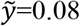 and 0.25 along the A-P axis. Position of the *otd* domain in mutants can be predicted straightforwardly by finding 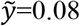 and 0.25 points in the corresponding mutant fate maps (Fig. 4e). These predictions are again in agreement with experiments^61^: decreasing Bcd dosage shifts both the anterior and posterior *otd* boundaries anteriorly, while lacking Tor makes the *otd* domain fail to retract from the anterior pole, and shifts its posterior boundary slightly forward. These observations in mutants are directly related to underlying regulations (as in Ref.^62^), that Bcd and Tor both activate *otd* in defining its posterior boundary, while at the same time both repress *otd* (probably through *huckebein*) in setting the anterior boundary (Fig. 4f). The gap gene regulation network seems to be organized in such a way to allow scaling.

### Dissecting the Contribution of Each Morphogen at Each Embryonic Position

In our framework, scaling originates from the insensitivity of decoder output to the correlated fluctuations in local morphogen levels. The forms of such correlated fluctuations are determined by the morphogen profiles. Therefore, if the morphogen profiles were perturbed, the alignment between the *y*-constant lines and the decoder’s decision boundaries would be affected and scaling would be as least partially destroyed. To provide quantitative test from this perspective, fate maps in mutant embryos with non-standard length (*L*≠1.0) are studied. These fate-maps are obtained by substituting the morphogen profiles given by Eq. 5 (with *L*≠1.0) into the above constructed phenomenological decoder.

Nearly all the decision planes have non-negligible projections alone the Bcd axis according to the scaling decoder, implying that Bcd contributes to patterning throughout the entire embryo. Therefore, missing Bcd destroys scaling completely in the prediction – the gap gene pattern changes greatly with *L* (Fig. 5a). More specifically, with decreasing embryo length, the *bcd*^*-*^ embryo is predicted to lose domain iv (*Kr*) and then v (*kni*). Amazingly, this phenomenon is observed in a recent experiment^25^ (see Fig. S7c for a detailed discussion).

**Fig. 5.**
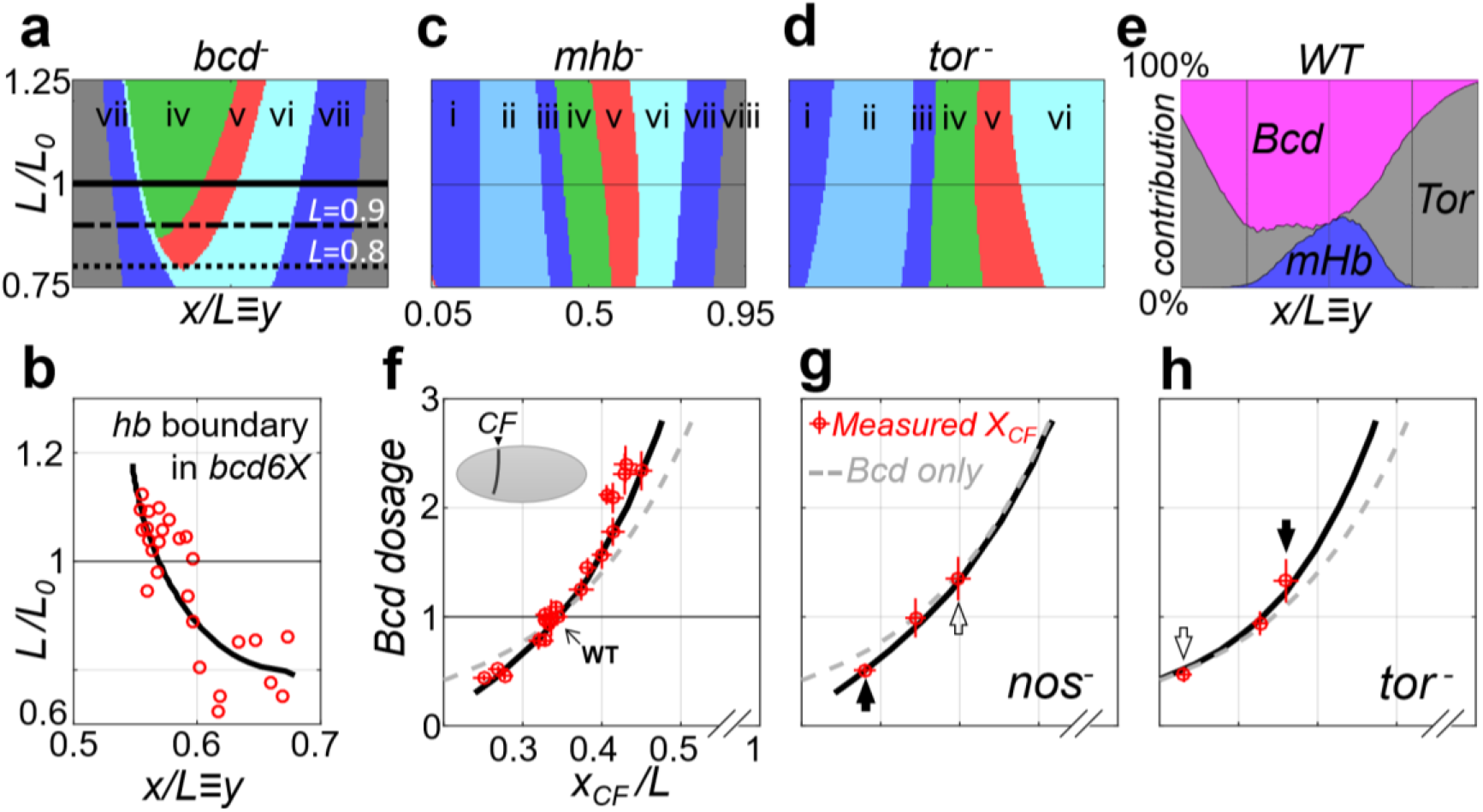
Different morphogens collaborate to achieve scaling in different regions. (**a**) Missing Bcd destroys scaling throughout the embryo. In *bcd*^*-*^, as *L* shrinks from 1.0 to 0.9 to 0.8 (solid, dashed and dotted lines), the *Kr* (iv) and *kni* (v) domains are predicted to disappear successively. (**b**) The effect of increasing Bcd dosage on scaling is accurately captured by the decoder. The position of predicted (black line) vs. measured (red circles)^25^ *hb* boundary (iii-iv boundary,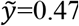) are shown. (**c**-**d**) In *mhb*^*-*^ and *tor*^*-*^, scaling is affected mostly in the middle and the two termini, respectively. (**e**) Relative contributions of the three morphogens to scaling across the embryo. (**f**) Shift of cephalic furrow (CF) under Bcd dosage change. Solid line is our prediction and red dots with error bars (standard deviation) are from experiment data^50^. Dashed grey line shows the position of the same Bcd concentration as CF in WT. (**g**-**h**) Shift of CF position in *nos*^*-*^ and *tor*^*-*^ backgrounds, respectively. The observed shifts match our predictions (solid line) well.

Furthermore, increasing Bcd dosage also affects the matching between the decoder decision planes and the *y*-constant lines, predicting that in embryos with additional *bcd* copies, scaling should also be affected. Experimentally, the mid-embryo *hb* boundary is indeed reported to be unscaled in shortened embryos when Bcd dosage is increased^25^ (Fig. 5b, red dots), which is accurately capture by our scaling decoder (Fig. 5b, black line. See also Supplemental text S8 for an analytical calculation).

In contrast to Bcd, at least some of the decision planes have almost zero projection alone the mHb or Tor axis, implying that missing mHb or Tor affects scaling in only part of the embryo (Fig. 5c, d). The relative contribution to patterning from each of the three morphogens (i.e., projection of each decision plane along the three axis) across the embryo can be quantitatively defined (Supplemental text S8) and are shown in Fig. 5e. Note that the morphogen mHb has long been considered dispensable or redundant, since losing mHb does not lead to any direct loss of the segment^44,45^. However, our results indicate that it in fact plays a crucial role for scaling in the abdominal region.

As a final example, we provide a quantitative explanation for the experimentally measured shift of the cephalic furrow (CF) under mutation of Nos or Tor plus Bcd dosage change. CF is the morphological boundary between head and thorax locating at *y*=0.344 in WT. It is a morphological trait much down-stream of the gap genes. But, as we believe that the scaling blueprint is established at the gap gene stage, we can regard CF as a hypothetical gap gene boundary at 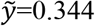 and predict its position *y*≡*X*_CF_ in various mutants. These predictions are compared with the measurements reported in^50^ (Fig. 5f-h).

When Bcd dosage is perturbed, the predicted *X*_CF_ matches well with the experimental measurement (Fig. 5f). It is clear that *X*_CF_ shows robustness against Bcd dosage change – its shift is always visibly *smaller* than that of the iso-concentration point of Bcd (Fig. 5f-h, gray dashed line). Such robustness of boundary positions has long been noted, and is frequently attributed to some unknown “self-correction” mechanism of the gap gene system^35,50,63^. In our framework, it is a direct reflection of the contributions from the other two morphogens.

Perhaps more convincing is the case when Bcd dosage is altered in the absence of mHb or Tor gradient. In *nos*^*-*^ mutant where mHb gradient is eliminated (Fig. 5g), with *increasing* Bcd dosage such robustness disappears, and *X*_CF_ follows the Bcd threshold exactly (marked by white arrow); while for *decreasing* Bcd dosage, such robustness still exists (black arrow). Similar but opposite phenomenon is seen in *tor*^-^ mutant (Fig. 5h). These phenomena are successfully captured by our model without any further tuning on parameters. Shift of CF in these cases can be easily understood from information provided in Fig. 5e – Bcd and Tor are the main contributing bi-gradient pair anterior to the CF, while the Bcd-mHb pair dominants the middle part behind CF. Therefore, positional robustness of the CF against Bcd dosage change has different origins when shifted anteriorly or posteriorly.

### The Scaling Decoder Can be Implemented with Gene Regulation

The discussions so far are quite general, independent of any details in the decoder’s biochemical implementation. In this section we provide evidence that the scaling decoder can be implemented by gene regulation and more specifically by the known gap gene network.

First, consider a toy model with a static anterior morphogen *B* (like Bcd) of fixed length constant and a single gap gene *H* whose initial condition *H*_init_ serves as the second posterior gradient (like mHb) (Fig. 6a). The desired output is to have *H* being activated in the anterior half with the boundary locating at *y*=0.5 regardless of the embryo length *L*. This requires that the morphogen levels marked by the black dots or empty triangles (Fig. 6a) should both lie on the *H* boundary. Let the regulation network be that of the inset in Fig. 6b. From a dynamical perspective, this network is well known to have bi-stability and Fig. 6b shows a typical bifurcation diagram. Within the bistable range, the *H-high/H-low* states are separated by a critical line (red squares), which is the achieved decision boundary between the two cell fates. If the activation strengths are properly tuned, this critical line can be adjusted to have the same slope as that of the desired decision boundary in Fig. 6c. Such first-order approximation is already good enough for ±20% variations in embryo length.

**Fig. 6.**
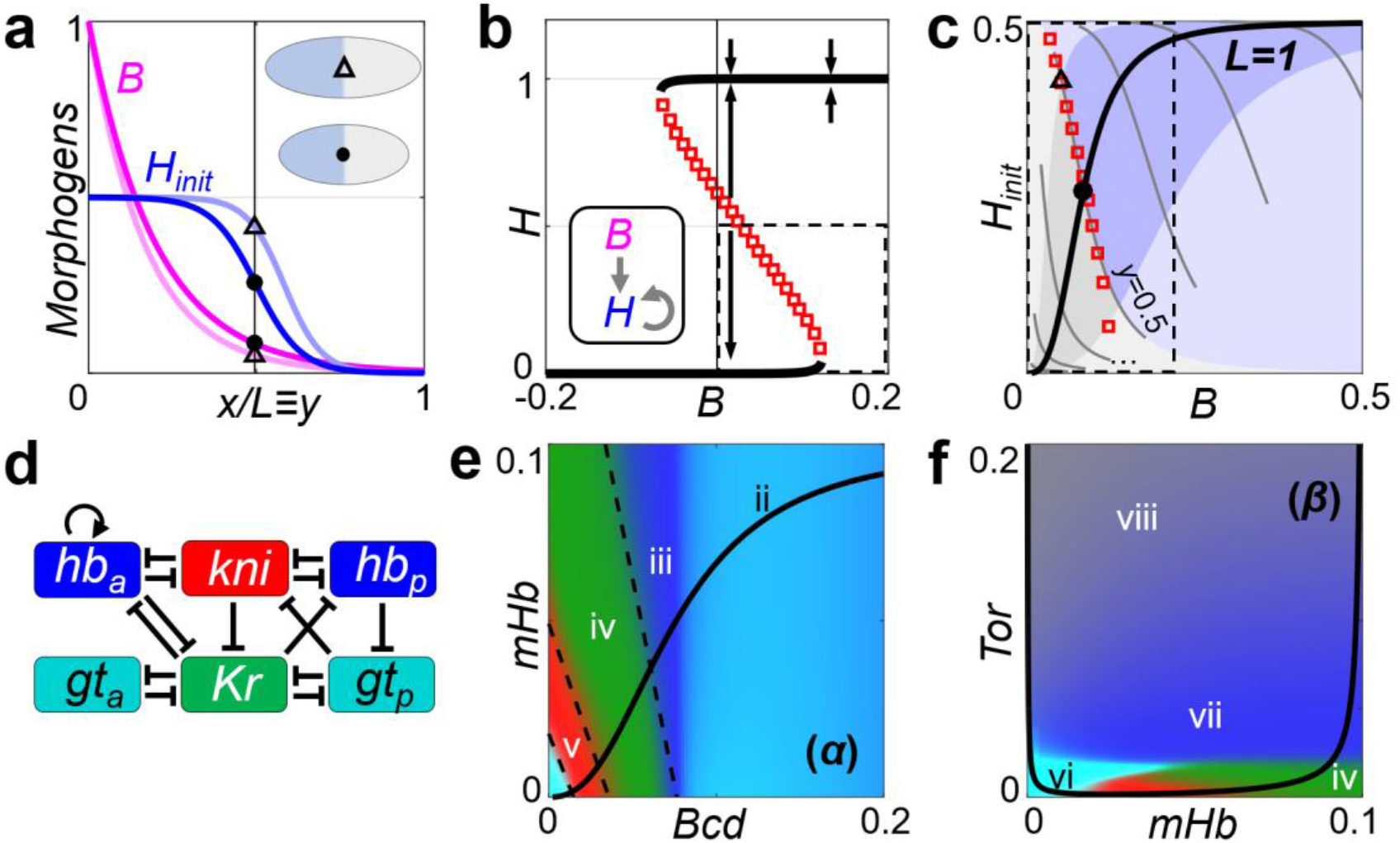
The scaling decoder can be implemented by gene regulation network. (**a**) The profiles of two morphogens in a toy model, shown for two embryos of standard size (darker lines) and larger size (lighter lines). (**b**) The bifurcation diagram for the network in the inset. A cell can reach one of the two bi-stable states depending on the values of the two morphogens, separated by the unstable manifold (red squares). This unstable manifold is the decision boundary separating the H-high and H-low fates. Its slope can be tuned by the activation strengths. (**c**) Within the realistic *L* range (±20%, highlighted region), The *y*=0.5 line is well approximated by the dynamical decision boundary in (**b**) (red squares). (**d**) The known gap gene network from Ref.^42^ (**e**-**f**) The same plots as Fig. 3A-B, but using an ODE model of (**d**) (see Supplemental text S11 and Fig. S11 for details). Dashed black lines are decision boundaries of the phenomenological decoder, same as Fig. 3a.

The gap gene network has more degrees of freedom and more scaling boundaries, and does not necessarily have to reach a dynamical attracting point. But the general idea is the same, that decision boundaries of the decoder correspond to boundaries between dynamical attracting basins. Based on the known gap gene network shown in Fig. 6d^42^, we construct a differential equation model (see Supplemental text S11 for details). If fitted to the WT data^27^ plus scaling requirement, the model can produce very similar decision boundaries as the phenomenological decoder (Fig. 6e, f), and hence correct predictions on mutants (Fig. S11).

## Discussion

Scale-invariant gene patterning can originate from an integrative decoding of non-scaling signals. In such case, the effective input-output structure of the decoder is largely dictated by the requirement of scaling. Consequently, the underlying gene regulation system evolves to have its macroscopic behavior approximating the ideal decoder geometry. All the intricate biochemical details seem to be hidden and largely irrelevant.

We provided strong evidence that scaling in the *Drosophila* gap gene expression pattern indeed emerge in this way. There are quantitative agreements between the predictions from the scaling decoder geometry and the experiments on nearly all maternal morphogen mutants. We then showed that the scaling requirement also contains rich information about the overall gene regulation logic and the way the morphogen signals are being integrated, all these inferred properties are consistent with the existing knowledge of the gap gene regulation network. The decoder was further demonstrated to be implementable by dynamical gene regulation models.

We note that the concept of the optimal decoder (i.e., decoder structure determined by its function) is not new even within the *Drosophila* community^28,40^. Our contribution here is that we have provided a unified understanding to scaling, mutants’ behavior and gene regulation in the *Drosophila* gap gene system using an “optimal decoder”. Importantly, we think that scaling is a central goal for the *Drosophila* decoder – as the morphogen profiles (e.g., Bcd) are already quite precise, their molecular noise should not be a major factor shaping the geometry of an optimal decoder. In Supplemental text S10 we demonstrate that a decoder optimally designed for correcting small and homogeneous morphogen noise is not optimal in the sense of scaling, nor does it explain the mutants’ behavior as satisfactorily as the one starting from scaling.

Finally, since our framework is rather general, the arguments presented in this paper can be readily applied to other systems to test if scaling is achieved through integrating non-scaling morphogens. In Supplemental text S13, we briefly discuss the situation of another long-germband insect *Megaselia abdita*.

## Methods

### Fitting the linear classification planes

The WT point cloud (Fig. 2d) used for fitting the linear classification planes are generated as follows. First, the A-P axis is discretized into 101 points *y*=0% to 100%. For each of the *y* position, we sample 400 embryo length values from the normal distribution *L*∼N(1, 0.1) and calculate the corresponding (Bcd, mHb, Tor) levels using Eq. 5. The three morphogen values are noted by ***m***=(*m*_1_, *m*_2_, *m*_3_) for convenience. Obviously, 0<*m*_*1,3*_<1 and 0<*m*_*2*_<mHb_0_ (mHb_0_=0.1 here).

Then, a Poisson noise is added to each *m*_*i*_ by assuming the actual number of molecules is a Poisson variable *n*_*i*_ with <*n*_*i*_>=*N*\**m*_*i*_, and the final morphogen value with noise is *n*_*i*_/*N*. The maximum number *N* controls the noise magnitude, and set to *N*=1000 throughout the main text. Note that we do not claim that the *actual* morphogen noise follows independently identically Poisson distribution. And *N* does *not* correspond directly to the number of molecules per nucleus. Instead, *N*=1000 is chosen to make the strength of the positional error of Bcd gradient close to that measured by Gregor, et. al.^51^ (about 1∼2% embryo length in the anterior half of embryo). After all, our model is not sensitive to the exact value of *N*, see Fig. S4e, f for the results for *N*=500 or 2000.

The entire point cloud in Fig. 2d thus consists of 101 subsets ***m***|_*y*_, each of them have 400 points. The 100 classification planes are located at *y*=0.5%, 1.5%, …, 99.5%, denoted as classifiers #1, …, #100. Since gap gene expression is affected by the dorsoventral system when being very close to embryo termini, only classifiers #6 to #95 are considered in the main text. Each of the planes should perform the *local* classification task of distinguishing adjacent ***m*** point sets. For example, the plane locates at *y*=2.5% should go through the *noise-free y*=2.5% point for *standard-sized* embryo ***m***|_*L*=1,*y*=2.5%_ by itself. And its orientation is such that it can best distinguishing the point sets **⋃**_*L*_ **⋃**_*y*={0%,1%,2%}_ ***m***|_*y*_ against **⋃**_*L*_ **⋃**_*y*={3%,4%,5%}_ ***m***|_*y*_. To find the best plane orientation numerically, we simply enumerate the Euler angles *θ* and *φ* of its normal vector at the resolution of 1° and find the one with the highest classification accuracy.

The noise due to finite sampling and discretizing *θ* and *φ* can be eliminated by averaging the classification plane orientations for 25 repeats of the above sampling and fitting steps.

### Predicting the fate-map function with the ensemble of linear classifiers

In principle, 100 well-separated planes should divide the morphogen space (the cube with 0<*m*_*1*_<1, 0<*m*_*2*_<0.1, 0<*m*_*3*_<1) into 101 slices, corresponding to 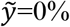 to 100%. If a query ***m*** point falls into the slice 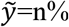, it locates on the posterior side of classification planes #1 through #n, and on the anterior side of planes #n+1 to #100. Therefore, the corresponding cell fate 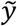 can be read out from the classification results of all the linear classification planes (Supplemental text S3).

In some situations, however, the 100 linear classifiers could have contradictory outputs. Say, a point may be classified to the posterior side by classifier #70 but to the anterior side by #30. We introduce a “posterior dominance rule” to tackle this difficulty. Anytime when this happens, output of the anterior classifier (#30 here) is always ignored. The reason for us to introduce the posterior dominance rule is simple and empirical – some anterior classification planes may intersect with the much more posterior region of the point cloud, vary far from where they were fitted (Fig. S3b). This posterior dominance rule works well, and finally yields all the results in Figs. 2-5.

## Data availability

All data generated or analyzed during this study are included in this published article and supplemental information.

## Acknowledgments

We thank Timothy Saunders for providing data for Fig. S7b, d, and e. This work was supported by the National Natural Science Foundation of China (12090053, 32088101).

