## Supplemental text for "Scaling dictates the decoder structure"

**Content.**

**Supplemental text.**

**S1.** Shape of the maternal morphogen profiles.

**S2.** Modelling length fluctuation and noisy morphogen gradients.

**S3.** Fitting the linear classification planes.

**S4.** Making predictions with the set of linear classifiers.

**S5.** The Bayesian decoder.

**S6.** Discussion on some of the predictions in Fig. 3d.

**S7.** *bcd*$^-$ and *bcd6X* embryos with reduced lengths.

**S8.** The three maternal gradients in *Drosophila* function as two bi-gradient pairs.

**S9.** Shift of the *eve* stripes under Bcd dosage change.

**S10.** Optimal decoder for noisy gradients without embryo length variation.

**S11.** Implementing the scaling decoder with a dynamical gene regulation model.

**S12.** Nos/mHb as the posterior gradient.

**S13.** Discussion on the long-germband insect *Megaselia abdita*.

**Supplemental figures.**

**Fig. S1.** The mHb profile can be fitted by a sigmoidal curve.

**Fig. S2.** Simulating noise in the Bcd gradient by a Poisson noise term.

**Fig. S3.** Predicting mutant fate-map with linear classifiers.

**Fig. S4.** Effects of the Bcd $\beta$ factor and noise amplitude on model performance.

**Fig. S5.** The Bayesian decoder.

**Fig. S6.** Outputs of the scaling decoder on mHb=0 and mHb=$\text{mHb}_0$ planes.

**Fig. S7.** Maternal morphogen mutant embryos with greatly changed length.

**Fig. S8.** The three maternal gradients function as two bi-gradient pairs.

**Fig. S9.** Shift of the *eve* stripes under Bcd dosage change.

**Fig. S10.** Optimal decoder for noisy gradients with or without length variation.

**Fig. S11.** The ODE-based gap gene regulation model.

**Fig. S12.** The scaling gene circuit model.

**Fig. S13.** Generating scaling *Drosophila* gap gene pattern with a more extended Bcd profile.

**Supplemental tables.**

**Table S1.** Peak and boundary positions extracted form maternal morphogen mutants.

**Table S2.** Parameters for the scaling ODE model.

**Table S3.** Parameters for the scaling gene circuit model.

**S1. Shape of the maternal morphogen profiles.**

**Bicoid (Bcd).** The Bcd concentration gradient is generated by diffusion form a localized source. At steady state, its profile should be exponential, which is fully consistent with experiments (therefore, we do not consider the possibility raised in Ref. [1]):

$$Bcd(x) = e^{-\frac{x}{\lambda_B}}.$$

Its absolute length constant $\lambda_B$ is fully determined by the diffusion constant $D$ and decay rate $\gamma$ ($\lambda_B = \sqrt{D/\gamma}$), independent of embryo length [2-5]. Reformulating the Bcd profile using the "relative" coordinate $y$, which is normalized by embryo length $y \equiv x/L$, yields:

$$Bcd(y) = e^{-\frac{yL}{\lambda_B}} = e^{-\frac{y}{\lambda_B/L}}.$$

For larger embryos ($L>1$), the length constant appears to be shortened in the normalized coordinate.

Throughout this paper, the length unit is chosen to be the length of a "standard size" embryo $L_0$ (~490 μm). Therefore, the position $x$, embryo length $L$, and $\lambda_B$, are dimensionless (normalized by $L_0$). Measured in this way, the Bcd length constant $\lambda_B$=0.165, according to a very carefully performed quantitative measurement [6].

Though the Bcd length constant $\lambda_B$ cannot scale with embryo length, a positive correlation between Bcd amplitude (absolute concentration at the anterior pole) and embryo length $L$ has been observed experimentally [5,7]. i.e., larger embryos tend to have higher overall Bcd dosage. To reflect this fact, an amplitude factor $L^\beta$ is introduced in the Bcd term. Taken together,

$$\mathrm{Bcd}(y, L) = L^\beta e^{-\frac{y}{\lambda_B/L}} \qquad (S1.1)$$

The exponent $\beta$ should lie between 2 and 3 according to a not very precise measurement [8]. Being somewhat conservative about this "amplitude correction" effect, we take $\beta$=2 throughout the main text. No matter if $\beta$=3 (Fig. S4d). Our model is not quite sensitive to the exact value of $\beta$.

This effect produces a "neutral point" $y=\beta\lambda_B$ with invariant Bcd concentration and is proposed by some authors that is may contribute directly to scaling of the gap genes. As mentioned in the main text, we don't agree with this explanation in general. However, in our framework, although the optimal scaling decoder can always be well-defined with or without this $L^\beta$ factor, the exact Bcd profile affects the exact orientations of the optimal decision planes. Therefore, to accurately describe the real situation in *Drosophila* (hence making correct predictions on mutants), this Bcd amplitude effect should not be ignored (Fig. S4b, c).

**Maternal Hb (mHb).** Like Bcd, the posterior gradient Nos should have an exponential profile. (No amplitude correction effects are reported experimentally, so the $L^\beta$ factor is not added).

$$Nos(y) = e^{-\frac{1-y}{\lambda_N/L}}$$

It is well known that Nos functions solely through repressing the maternal component of the gap-gene protein Hb (mHb) in posterior half of the embryo [9-11]. Therefore, the "immediate" posterior morphogen should be mHb instead of Nos. If we assume that mHb level is dictated by Nos through an inhibitive Hill function,

$$mHb(y) = \frac{mHb_0}{1 + \left(\frac{Nos(y)}{K}\right)^n}$$

then the mHb gradient takes a sigmoidal shape:

$$mHb(y) = \frac{mHb_0}{1 + e^{\alpha L(y-1+(1-\lambda_H)/L)}} \qquad (S1.2)$$

Where $\alpha = n/\lambda_N$, $\lambda_H = 1 + \lambda_N \ln K$. Although the raw parameters ($\lambda_N$, $K$, and $n$) are unknown, measured mHb profiles (Hb protein profile in n.c.12 embryos, from the *FlyEX* database [12,13]) can be well fitted with this sigmoidal curve (with $\alpha$=15 and $\lambda_H$=0.425). See Fig. S1 for the fitting. Note that in this paper we normalize the value of Hb (no matter maternal or zygotic) according to its maxima at n.c.14, so mHb has an amplitude coefficient $mHb_0$=0.1.

**Torso (Tor).** The activity of Tor is induced at both terminus of the embryo by its ligand in the perivitelline space [14,15]. Tor transduces the activation signal into the syncytial embryo by phosphorylating ERK. Phosphorylated ERK (dpERK) diffuses in the cytoplasm, trapped by the nucleus, and dephosphorylated (i.e. “degraded”) inside the nucleus [16]. This is a similar picture of the “localized synthesis, diffusion, and decay” model of Bcd and Nos. So, it is reasonable to assume that the activity of Tor has an exponential profile as well.

$$Tor(y) = e^{-\frac{y}{\lambda_T/L}} + e^{-\frac{1-y}{\lambda_T/L}} \qquad (S1.3)$$

Quantitative measurements on dpERK indeed show double-exponential profiles (when projected onto the one-dimensional anterior posterior axis) [16,17]. In this paper we use an estimated value $\lambda_T$=0.07 for its length constant.

These equations give Eqn. 5a-c in the main text.

Precise measurements on the scaling property of mHb and Tor is still lacking and very difficult to perform. Our assumption that they should both be unscaled with embryo length is the minimal assumption. This minimal assumption is consistent with the known mechanisms through which the gradients are established, and is also supported by the case studied in Fig. 5a. See SI-7 for detail.

**S2. Modelling length fluctuation and noisy morphogen gradients**

The WT point cloud used for fitting the linear classification planes are defined as follows. First, the A-P axis is discretized into 101 points $y$=0% to 100%. For each of the $y$ position, we sample 400 embryo length values from the normal distribution $L \sim N(1, 0.1)$ and calculate the corresponding noise-free (Bcd, mHb, Tor) levels using Eqn. 5. This three noise-free values are noted by $\boldsymbol{m}=(m_1, m_2, m_3)$ for convenience. Obviously, $0<m_{1,3}<1$ and $0<m_2<0.1$. This give the 2-d "WT manifold" in Fig. 2c.

The fact that embryo length $L$ only fluctuates within a limited range is important. Outside certain $L$ range, the decoder behavior should not be subjected to selection pressure since embryo size hardly fluctuate that much under natural conditions. Thus, in the strictest sense, only within the region covered by *realistic* WT embryos, the effective input-output relation of the decoder should follow that dictated by scaling.

Secondly, a Poisson noise is added to each morphogen value $m_i$ by assuming the actual number of molecules is a Poisson variable $n_i$ with $<n_i>=N*m_i$, and the final (normalized) morphogen level with noise is $m_i=n_i/N$. The (hypothetical) maximum molecule number $N$ controls the noise magnitude. We set $N$=1000 throughout the main text. From a theoretical perspective, the noise terms turn the 2-dinemsional WT manifold (Fig. 2c) into a 3-dimensional WT point cloud (Fig. 2d).

Note that $N$=1000 does not correspond to the number of molecules per nucleus (which result in the intrinsic noise). Instead, $N$=1000 is chosen to make the positional error of modeled Bcd gradient close to that measured by Ref.[18], which included both intrinsic and extrinsic noises.

To be specific, positional noise (standard deviation $\sigma$ of position $y$) of the Bcd gradient in standard-sized ($L$=1) embryos can be expressed as

$$\sigma_y \left|\frac{d\bar{N}_{Bcd}}{dy}\right| = \sigma_{N_{Bcd}} \qquad (S2.1)$$

For Poisson distribution: $\left(\sigma_{N_{Bcd}}\right)^2 = \bar{N}_{Bcd} = Ne^{-y/\lambda}$. Thus:

$$\sigma_y = \frac{\lambda}{\sqrt{N}} e^{-\frac{y}{2\lambda}} \qquad (S2.2)$$

Substituting $\lambda$=0.165 and N=1000 into this equation, gives the modeled positional noise as the black curve in Fig. S2, overlapped on the experiment results of (Gregor, et. al. 2007).

The measured results on Bcd noise should be reliable in the region 0.2<y<0.6, since it is far enough from Bcd mRNA is distribution, also, Bcd protein level here is high enough to be safe from experimental detection limit. In this region 0.2<y<0.6, strengths of Bcd noise introduced by our Poisson term are close to the measured values. (Our model is not quite sensitive to the exact value of *N*, see Fig. S4 E-F for the results when *N*=500 or 2000.)

Note that "Bcd noise" here should stand for the measured embryo-to-embryo fluctuation in an ensemble of standard-sized (*L*=1) embryos. i.e., it accounts for both intrinsic noise (finite number of Bcd protein molecules per nucleus), and extrinsic noise (e.g., embryo-to-embryo variability in overall Bcd amplitude) except for length variation. Theoretically, intrinsic and extrinsic noises are different, in that extrinsic noise is correlated for nucleus belonging to the same embryo. However, since our decoder works in a *spatially decoupled* manner, it cannot distinguish whether two different (Bcd, mHb, Tor) points come from the same embryo or not. In other words, only the overall strength of fluctuation matters, no matter the fluctuation comes from intrinsic molecular noise or embryo-to-embryo variation. The decoder deals with all nucleus in all embryos of the *Drosophila* species simultaneously. Therefore, intrinsic and extrinsic noises are not treated separately here.

**S3. Fitting the linear classification planes.**

The entire point cloud in Fig. 2d consists of 101 subsets $\boldsymbol{m}|_y$, each of them have 400 points. The 100 classification planes locate at *y*=0.5%, 1.5%, …, 99.5%, numbered as classifiers #1, …, #100 (and we only consider #6 to #95 in the main text). Each of the planes should perform the *local* classification task of distinguishing ***m*** points belongs to adjacent *y*'s.

For example, the plane #3 locating at *y*=2.5% should first go through the ***m*** point representing the *noise-free* morphogen levels at *y*=2.5% in *standard-sized* WT by itself. Secondly, the plane orientation is defined by that can best distinguishing the point classes $\bigcup_{y=\{0\%,1\%,2\%\}}\boldsymbol{m}|_y$ against $\bigcup_{y=\{3\%,4\%,5\%\}}\boldsymbol{m}|_y$. To find the best-fit plane orientation numerically, we simply enumerate the Euler angles $\theta$ and $\varphi$ of its normal vector at the resolution of 1° and find the one with the highest classification accuracy.

The noise due to finite sampling and discretizing $\theta$ and $\varphi$ are eliminated by averaging the classification plane orientations for 25 repeats of the above sampling and fitting steps.

**S4. Making predictions with the set of linear classifiers.**

The portion of (Bcd, mHb, Tor) space where the decoder output is directly dictated by scaling (that is, the region covered by the WT point cloud) does not include all the situations in the morphogen mutant embryos. Extrapolations are therefore needed for making predictions in general. Fortunately, for the *Drosophila* case most mutants of interest lie not far away from the WT point cloud. Thus extrapolations could make sense here.

We think the most simple and natural assumption is to extrapolate linearly with the classification planes defined above.

Firstly, we know from Fig. 6c that the decision boundaries realized by gene interaction network can only follow the scaling requirements to the linear order in general, thus starting only from scaling we simply have no information about possible higher-order features.

Secondly, the unstable manifold of a bi-stable diagram tends not to have large curvature except in the neighborhood of a critical point (Fig. 6b). Although the real gap gene network is not a simple bi-stable system, we think this intuition should still hold.

Finally, in Fig. 6d-f we presented a differential-equation-based gene regulation model using the known gap gene interaction network. This kind of model indeed extrapolate in a very much linear way.

We next describe how the linear extrapolations are carried out precisely. Obviously, 100 well-separated planes should divide the morphogen space (the cube with $0<m_1<1$, $0<m_2<0.1$, $0<m_3<1$) into 101 slices, corresponding to $\tilde{y}$=0% to 100%. If a query $\boldsymbol{m}$

point falls into the slice $\tilde{y}$=n%, it locates on the posterior side of classification planes #1 through #n, and on the anterior side of planes #n+1 to #100. Therefore, the corresponding cell fate $\tilde{y}$ can be read out from the classification results of all the linear classification planes. Consider the $y$=0.4 point in an $L$=1 $bcd^{-}tor^{-}$ embryo, whose $\boldsymbol{m}$=(0,0.059,0) (according to Eqn. 5, shown as black cross in Fig. S3a). This point lies in the $\tilde{y}$=0.56 slice, i.e., between the classification planes #55 and #56.

In some other situations, however, the 100 linear classifiers could have contradictory outputs. Say, the point (0.004, 0, 0.24) are classified to the posterior side by classifier #70 but to the anterior side by #30. We introduce a “posterior dominance rule” to tackle this difficulty. Anytime when this happens, output of the anterior classifier (#30 here) is always ignored. The reason for us to introduce the posterior dominance rule is simple – some anterior classification planes may intersect with the much more posterior region of the point cloud, vary far from where they were fitted (Fig. S3b). This posterior dominance rule works well, yielding the results in Figs. 2-5.

To be more precise how the fate map curves in Fig. 3 are obtained, we explain here that which “fate slice” a query point should fall is determined by analyzing the classification outputs by all the 100 classification planes. In the ideal case, $\tilde{y}$=n% means that classifiers #1 through #n output “posterior”, and classifiers #n+1 to #100 output “anterior”. Graphically, the decoder outputs are recorded by a column of pixels, with grey stands for “anterior” and white stands for “posterior”, and the fate map in Fig. 3 is then given by extracting the grey-white boundary of Fig. S3c.

Also note that those classifier outputs ignored by the posterior dominance rule are shown in lighter color in Fig. S3c. Graphically, this rule is equivalent to wiping out all the grey pixels once there exists a white pixel above them. The “posterior dominance rule” is a quite rough rule after all. Sometimes it leads to artifacts near the embryo terminus. For example, the fate map of *bcd1X* (and $vas^{-}exu^{-}bcd6X$) in Fig. 3 shows abrupt jumps near the anterior (and posterior) end. However, this kind of error do not affect prediction in most cases.

**S5. The Bayesian decoder.**

With our settings, given position $y$ and embryo length $L$, the morphogen levels $\boldsymbol{M}$ satisfy the Poisson distribution:

$$p(M_i|y,L) = \frac{\left(Nm_i(y,L)\right)^{M_i}}{M_i!} e^{-Nm_i(y,L)} \quad (i = 1,2,3),$$

where $m$ is the *noise-free* morphogen profile defined in Eqn. 5 in main text, and $N$ is the "effective maximal molecule number" as described above. Also, noise on different morphogens are assumed to be independent:

$$p(\boldsymbol{M}|y) = \int dL\, \rho(L) \prod_{i=1}^{3} p(M_i|y,L),$$

where the embryo length follows the Gaussian distribution $L \sim N(\mu=1, \sigma=0.1)$

$$\rho(L) = \frac{1}{\sqrt{2\pi}\sigma} e^{-\frac{(L-1)^2}{2\sigma^2}}$$

Inferring $y$ from the morphogen levels $\boldsymbol{M}$ can be performed using the Bayes formula

$$p(y|\boldsymbol{M}) = \frac{p(\boldsymbol{M}|y)\, p(y)}{p(\boldsymbol{M})}$$

Since $y$ is uniformly sampled from 0 to 1, the Bayesian decoder becomes a maximal likelihood decoder.

$$\underset{y}{\operatorname{argmax}}\, p(y|\boldsymbol{M}) = \underset{y}{\operatorname{argmax}}\, p(\boldsymbol{M}|y)$$

Decoding results for the WT point cloud of Fig. 2d by the Bayesian decoder is shown in Fig. S5a. Although arbitrary decision boundary geometry is allowed by the Bayesian decoder, its Root Mean Squared Error (RMSE) is even larger than the linear decoder presented in the main text (Fig. S5b). This is conceivable, as maximization of posterior likelihood could lead to minimal regression error only when the decoding error is Gaussian, which not satisfied here. Therefore, we claimed in the main text that the remaining classification errors of the linear decoder should due to the morphogen noise rather than nonlinearity in classifier geometry. This can be visualized by choosing a tangential view of the classification planes in Fig. 2D-E (Fig. S5d).

In comparison with the Bayesian decoder, there are some additional arguments on our linear extrapolation hypothesis. As expected, decision boundaries of the Bayesian decoder are effectively linear within the WT point cloud (Fig. S5c, on the alpha plane). However, situations outside the WT point cloud are quite different: There, outputs of the Bayesian decoder are determined by extremely improbable cases, say very large $L$ variation or very large fluctuation in morphogen level, which are basically irrelevant to realistic embryos. While the linear classifier emphasizes more on simplicity of the decision boundary geometry, and turned out to be a better way of extrapolation.

**S6. Discussion on some of the predictions in Fig. 3d.**

**(*mhb*$^-$)**. Among the mutant cases in Fig.3d, we do not have quantitative data for *mhb*$^-$, but we are quite confident that this prediction is correct. It has long been known that although *nos*$^-$ embryos (where mHb is uniformly high) lack all abdominal segments, it can be largely rescued by further eliminating mHb completely. The *nos*$^-$*mhb*$^-$ double mutation embryo is viable and has basically normal morphology [9,10]. This observation led people to discover that for WT the only function of Nos is to inhibit mHb, but it also shows that even if the morphogen mHb does not exist at all, the embryo should not miss any segment. Our prediction in *mhb*$^-$ matches this observation. The predicted fate map is almost along the diagonal albeit slight distortions. Fig. S6 provides a graphical visualization on the case of *nos*$^-$ and *mhb*$^-$.

**(*vas*$^-$*exu*$^-$*bcd6X*)**. The *vas*$^-$*exu*$^-$*bcd6X* embryo lacks *nos* and has flattened and overall increased Bcd gradient [19], and the measured *bcd* profile is used for this prediction. This mutant display mirrored head structures near the posterior pole. The yellow squares marked in Fig. 3d *vas*$^-$*exu*$^-$*bcd6X* panel represent the expression peaks and boundaries of the head gap genes *otd*, *btd*, and *ems*, measured by Ref.[19].

**(*bcd1X* and *bcd4X*)**. *bcd1X* or *bcd4X* here means the situations where Bcd dosage is halved or doubled exactly, not the actual *bcd* copy number. Hb boundary in *bcd1X* or *bcd4X* is predicted to shift by -7.3% or +9.1% by our scaling decoder, and the experimentally measured shifts are -6.4% and +9.4% according to Ref. [6].

**S7. *bcd*$^{-}$ and *bcd6X* embryos with reduced lengths.**

The case of *bcd*$^{-}$ has been discussed in the main text. Our model predicts that domains iv and v should disappear successively (main text Fig. 5a), this is visualized graphically in Fig. S6a as the *bcd*$^{-}$ curve losses contact with the green and red regions successively as *L* shrinks.

This prediction is consistent with the experiments of Ref. [7], where *bcd*$^{-}$ embryos with greatly reduced embryo lengths are obtained by *fat2RNAi*. We replot the experiment data (Fig. 6B of Ref. [7]) here for a quantitative comparison (Fig. S7b, the narrow anterior *gt* domain is ignored). In this figure, each horizontal bar represents a measured gap gene expression domain in a *bcd*$^{-}$ or *bcd*$^{-}$*fat2RNAi* embryo. Its position in the vertical direction represents its embryo length. With decreasing length, the *kni* and *gt* domains shifts anteriorly in the normalized coordinate *y*. Also, *Kr* and *kni* domains disappears successively, leaving a widened *gt* domain in the middle.

To allow quantitative comparisons, it worth noting that the length of the fixed embryos used in the immunostaining experiments is in general shrinked. We still have some difficulties in figuring out all the complexities and experimental details regarding the shrinking ratio (i.e., the ratio of the embryo length after fixation and immunostaining to that of the same embryo when still alive). Therefore, the shrinking ratio is *assumed* to be 90% here, which makes the experiments and our model prediction overlaps satisfactorily (Fig. S7c).

This result directly supports our basic assumption on the maternal morphogens, that mHb and Tor gradients should be unscaled with *L*. In this picture, both posterior Tor and mHb gradients measure the absolute distance from the posterior pole, and the gap gene domains v-viii should simply follow these fixed distances and move anteriorly in relative coordinate as *L* shrinks. Similarly, the anterior bands (reversed domains vii and viii) are kept at fixed distances from the anterior pole by reading the anterior Tor gradient. In between is the *Kr* domain, which shrinks with *L*.

Using the *fat2RNAi* technique, Ref. [7] also studied the gap gene patterns in *bcd6X* embryos under length change. Note that with 6 copies of *bcd*, the resulting Bcd protein dosage was actually only be approximately doubled [20], rather than multiplied by 3. This knowledge is consistent with the position of cephalic furrow reported in [7], that CF locates at $y$=0.42 in the *bcd6X* embryos, very close to those Bcd dosage ≈ 2.2 embryos reported by Ref. [6]. So, Bcd dosage is set to 2.2 in our model to simulate these *bcd6X* embryos. Scaling is not preserved in these embryos – that the gap gene domain boundaries shift significantly when *L* changes (Fig. S7d). This is the expected result, as according to our basic assumption, scaling stems from cancelation of the first-order effects of morphogen level difference due to a change of *L*. Those first-order derivatives are different for the altered zeroth-order profiles, thereby failed to cancel each other out.

In Fig. S7d, the measured boundary positions by [7] are shown as dots, while our predictions are the solid lines (no “shrinking ratio” is assumed here). Errors between predictions and the experiments are generally acceptable, except for some of the boundaries – namely, both boundaries of the *kni* domain, and the anterior boundaries of the posterior *gt* and *hb* domains. Note that these errors could be corrected by more careful data processing. And the observation that the *kni* and posterior *gt* domains may disappear in short enough embryos [7] does not seem to be captured by our extrapolation based predictions.

Especially, shift of the mid-embryo *hb* domain can be studied analytically, because the effect of Tor is nearly negligible in the central region. Profiles of Bcd and mHb when Bcd dosage is 2.2 are as follows.

$$\begin{cases} \mathrm{Bcd}(y', L') = 2.2\, L'^{\beta} e^{-y'L'/\lambda_B} \\ \mathrm{mHb}(y', L') = \mathrm{mHb}_0\left(1 + e^{\alpha L'(y'-1+(1-\lambda_H)/L')}\right)^{-1} \end{cases} \qquad (S7.1)$$

Assume that there exists a WT embryo of length $L$ with perfectly scaling gap gene pattern, in which the $y$=0.473 position (*hb* boundary) has identical Bcd and mHb values as the above equation. i.e.,

$$\begin{cases} L^{\beta} e^{-yL/\lambda_B} = 2.2\, L'^{\beta} e^{-y'L'/\lambda_B} \\ \mathrm{mHb}_0\left(1 + e^{\alpha L(y-1+(1-\lambda_H)/L)}\right)^{-1} = \mathrm{mHb}_0\left(1 + e^{\alpha L'(y'-1+(1-\lambda_H)/L')}\right)^{-1} \end{cases}$$

Relationship between *y'* and *L'* can be easily solved by eliminating the unknown *L*:

$$L' = \lambda_B \frac{\ln 2.2 - \beta \ln \frac{1-y'}{1-y}}{1 - \frac{1-y'}{1-y}} \qquad (S7.2)$$

This analytical result is compared with experiments in Fig. S7e. Note that this prediction not even depends on the precise profile of mHb.

Similar to Fig. 5a-d, we present predicted gap gene domain positions of many other maternal morphogen mutants with greatly changed embryo lengths in Fig. S7f. Some of them may be tested by experiments in the future.

**S8. The three maternal gradients in *Drosophila* function as two bi-gradient pairs.** The "tri-gradient system" of Bcd, mHb, and Tor can largely be decomposed into two parallel bi-gradient systems: Bcd&mHb in the middle part, and Bcd&Tor near both termini.

The contribution of each morphogen to scaling can be characterized as follows. From the results shown in Fig. 5a, c, d, we can calculate the derivative of the position $y$ where a certain cell fate $\tilde{y}$ appears with respect to embryo length $L$ (evaluated at $L$=1.0), defined as the size sensitivity $S_L$.

$$S_L \equiv \lim_{\Delta L \to 0} \left| \frac{\Delta y}{\Delta L} \right| \qquad (S8.1)$$

With the fitted decoder classification planes, $S_L$ can be computed as follows. Let (Bcd($y$), mHb($y$), Tor($y$)) be the morphogen levels at $y$ in an embryo (WT or maternal morphogen mutant), the corresponding cell fate is $\tilde{y}(y)$. Since we would like to track the position where $\tilde{y}$ is fixed, the total shift in the (Bcd, mHb, Tor) space caused by $\Delta L$ and $\Delta y$ should be perpendicular to the normal vector $\boldsymbol{K}$ of the local decision plane:

$$\left(\Delta L \left(\frac{\partial Bcd}{\partial L}, \frac{\partial mHb}{\partial L}, \frac{\partial Tor}{\partial L}\right) + \Delta y \left(\frac{\partial Bcd}{\partial y}, \frac{\partial mHb}{\partial y}, \frac{\partial Tor}{\partial y}\right)\right) \cdot \boldsymbol{K}(\tilde{y}) = 0 \qquad (S8.2)$$

$S_L = \Delta y / \Delta L$ can thus be obtained.

$S_L$ values for WT are always below 0.1 (black curve in Fig. S8a-c); so, even standard deviation of $L$ is 10%, positional error introduced by imperfect scaling should be less than 1% (consistent with Fig. 2f). In *bcd*$^-$, patterns follow the fixed absolute distances to both termini (Fig. S8a, the dashed grey lines). When the Tor gradient is absent (Fig. S8c), $S_L$ increases significantly near both termini compared with WT, following that dictated by Bcd alone (dashed grey line); while in the middle part (*x/L* between 0.4 to 0.6) $S_L$ is still close to zero due to the remaining mHb gradient. The situation is similar when the mHb gradient is lost *(mhb*$^-$ or *nos*$^-$, Fig. S8b).

Taken together, these model predictions point to an intuitive interpretation to our phenomenological decoder. First, Bcd plays a central role throughout the entire embryo. Second, the gradient of mHb works together with Bcd as a pair of “bi-gradient” morphogens in the middle part ($y$ = 0.25 to 0.75), while Bcd and Tor works together near both ends (0 to 0.35, and 0.65 to 1.0).

This feature can also be illustrated from another perspective. We can directly calculate the contribution of each morphogen in discriminating adjacent points $y - \delta y/2$ and $y + \delta y/2$ in the WT embryo (for WT $y \equiv \tilde{y}$), i.e. specifying the cell fates in WT. The linear classifier sitting at position $y$ works by computing the sign of the inner product:

$$Z(y') \equiv \big(Bcd(y') - Bcd(y), mHb(y') - mHb(y), Tor(y') - Tor(y)\big) \cdot \boldsymbol{K}(y)$$

Z values for adjacent $y'$ points should differ by

$$Z(y + \delta y/2) - Z(y - \delta y/2) = \left(\frac{\partial Bcd}{\partial y}, \frac{\partial mHb}{\partial y}, \frac{\partial Tor}{\partial y}\right) \cdot \boldsymbol{K}(y)\delta y.$$

The contribution of Bcd, for example, is simply defined as the contribution of the Bcd term in this inner product:

$$c_{Bcd} = \frac{\frac{\partial Bcd}{\partial y} \cdot K_1(y)}{\left(\frac{\partial Bcd}{\partial y}, \frac{\partial mHb}{\partial y}, \frac{\partial Tor}{\partial y}\right) \cdot \boldsymbol{K}(y)} \qquad \text{(S8.3)}$$

Values of $c_{Bcd}$, $c_{mHb}$ and $c_{Tor}$ along the A-P axis are shown in Fig. 5e. Obviously, these three $c$ terms should add up to 1. The regions in which mHb and Tor play a role is clearly shown.

Note that we can define the Bcd dosage sensitivity $S_{\mathrm{Bcd}}$ in a similar manner. $S_{\mathrm{Bcd}}$ describes the shift $\Delta y$ of certain cell fate $\tilde{y}$ upon an infinite small change of Bcd dosage (by a factor of 1+ε, thus the Bcd exponential profile is effectively shifted by $\lambda_B \varepsilon$). $S_{Bcd} \equiv \Delta y/\lambda_B \varepsilon$. $\Delta y$ here is determined by

$$\left(\varepsilon(Bcd,0,0)+\Delta y\left(\frac{\partial Bcd}{\partial y},\frac{\partial mHb}{\partial y},\frac{\partial Tor}{\partial y}\right)\right)\cdot \boldsymbol{K}(\tilde{y})=0\,.$$

Since Bcd has an exponential profile, working out the formula yields that $S_{\mathrm{Bcd}}$ is in fact the same quantity as $c_{\mathrm{Bcd}}$ defined above.

An interesting point is that morphogen contributions are different for WT embryos of different lengths – larger embryos seem to rely more on mHb (Fig. S8d). This may be a testable prediction for future experiments – the same perturbation in mHb (or Nos) should introduce larger pattern shift (or more sever segment defects) for larger embryos (Fig. S8e).

Also note that Bcd functions throughout the entire embryo is also feasible biochemically. There should be still around 100 Bcd molecules per nucleus (concentration on the order of nM) even at the most posterior nucleus (estimated by the measurements of [18], with GPF maturation effect corrected following Ref.[6]).

**S9. Shift of the *even-skipped* (*eve*) stripes under Bcd dosage change.**

Shift of the cephalic furrow (CF, corresponds to the fate $\tilde{y}$=0.344) in response to Bcd dosage change (but embryo length is fixed to $L$=1.0) is discussed in the main text. This prediction can obviously be generalized to other "marks" on the fate map. For example, the seven *eve* stripes at $\tilde{y}$=(0.353, 0.435, 0.505, 0.56, 0.62, 0.675, 0.75) according to the measurements of [21]. Fig. S9a presents the predictions of the *eve* stripe positions under Bcd dosage change with or without the presence of the other maternal gradients (mHb and/or Tor). The intuitive explanation raised in Section S8 is again reflected in these results – that Bcd dosage robustness in the middle/terminal part depends on mHb/Tor (arrow heads). Note that since the 5th strip of *eve* locates at the posterior "boulder" between the region mainly governed by Bcd-mHb and that by Bcd-Tor, it should have exactly the same behaviors as CF discussed in Fig. 5f-h (which locates at the anterior "boulder" of this kind). This may be tested by future experimental studies.

Our predictions on *bcdnX* without further mutating mHb or Tor can be compared to the measurements of [22]. Their measurements on the *eve* stripes were carried out at some different developmental timepoint (and the embryo orientations are not well-controlled). So, we use *their* measured WT positions $\tilde{y}$=(0.33, 0.415, 0.5, 0.565, 0.635, 0.7, 0.78) to predict their results on *BcdnX* (Fig. S9b). The Bcd dosage of their *bcd4X* embryos is assumed to be 1.7 times of the WT since no exact values were provided.

**S10. Optimal decoder for noisy gradients without embryo length variation.**

Given the distribution of input signal, the decoder structure can be properly shaped to best utilize the available information, hence minimizing output error. This optimal decoding idea is commonly used in biological scenarios, ranging from neural sciences to developmental biology [23]. However, to get meaningful results via this approach, there should be much insight about the exact form of input fluctuations that the decoder really cares about – although minimizing output noise is always "a good thing", in many real-world situations, the decoder structure is shaped mainly by the requirements of other (more important) functions, not just by the simplest form of input noise alone.

In the case of *Drosophila* A-P patterning, we have considered two types of input fluctuations: noisy morphogen gradients and fluctuation in embryo length. We reason that the real-world *Drosophila* decoder must deal with both kinds of noise (especially the latter, i.e., scaling), since the decoder should function in all cells in all embryos of different lengths. Therefore, a decoder optimally designed only for noise attenuation in $L$=1 embryos is actually *not* optimal, by definition, for the *Drosophila* species in natural environment. In this section, we discuss the difference between the full scaling decoder (that in the main text, which takes into consideration both types of noise) and the optimal noise attenuation decoder where fluctuation in $L$ is not considered. By these analyses, we emphasize the importance of having the *correct* form of input noise in obtaining the *correct* optimal decoder structure.

The first situation studied here is called "noise-only decoder", where the Poisson noise term on morphogen gradients is the same as in Fig. 2d, but $L$ is now fixed to be 1 instead of being a random variable. The second model is a more drastic simplification, that morphogen noise is assumed to be vanishingly small. In this limit, each $\boldsymbol{m}(y)$ point is blurred into a small 3-d sphere, therefore, the optimal decision plane separating adjacent “spheres” is just the normal plane of $\boldsymbol{m}(y)$. We call it the “null model”, as it represents the most naïve form of “optimal” decoder one may imagine. Systematic investigation of the “noise-only decoder” and the “null model”, in comparison with the full “scaling decoder” in the main text, is summarized in Fig. S10.

First, the noise-only decoder and the null model *are* different from the optimal scaling decoder. This can be seen by direct visualization of the decision planes (Fig. S10a, e, i), which is further highlighted by the different extrapolations on the Bcd-mHb plane (Fig. S10b, f, j). Some of the differences are marked by arrowheads. Also, the morphogen contribution plots are different in three models (Fig. S10c, g, k). In Fig. S10d, h, the decoding result of these two other models on the full WT point cloud (that of Fig. 2d in the main text) are shown, scaling errors are in general larger compared to the scaling decoder (Fig. S10L). Therefore, the noise-only decoder and the null model are no longer optimal if fluctuations in embryo length are considered.

These structural differences are significant, since they lead to observable differences in mutant predictions (Fig. S10m, n, o). Predictions of our scaling decoder (panel o) match experiments the best, while the noise-only decoder or the null model has prediction error typically on the order of >5% embryo length (marked by arrowheads).

As an example, consider the vi-vii boundary (*gt-hb* boundary). In the scaling decoder, Bcd plays a non-negligible role near the posterior pole to allow for scaling (Fig. S10k). Graphically, the vi-vii boundary is inclined up (Fig. S10i, arrow head). This feature is supported by experimental measurements, that the posterior *hb* domain expands anteriorly in $bcd^{-}$ (Fig. 4b, Fig. S10r). However, this is not captured by the null model (Fig. S10q), nor the noise-only decoder (Fig. S10p), where this boundary is nearly parallel to the Bcd-mHb plane (Fig. S10a, e, arrowheads). In this example, the scaling decoder *is* the best description of *Drosophila* gap gene system among the three, as it predicts the mutant situation the most precisely.

Fig. S10s&t is in parallel with Fig. 5a of the main text, albeit for the noise-only and null models. Especially, the null model predicts that *Kr* and *kni* domains should disappear at nearly the same *L*, which contradicts experiments [7] qualitatively. Fig. S10 u&v is about the issue of cephalic furrow, in parallel with the main text Fig. 5f-h. Admittedly, the noise-only decoder predicts the CF position as satisfactorily as the scaling decoder. But the null model failed to predict the robustness of $X_{CF}$ with respect to Bcd dosage change (Fig. S10v), which is the most important feature here.

In summary, the analyses above demonstrate that scaling has indeed introduced some additional (and correct) constraints on the decoder structure, beyond merely decoding noisy gradients at fixed embryo length. The gap gene system seems to be "optimized" for both the kinds of fluctuations: noisy morphogen gradients and that in embryo length.

**S11. Implementing the scaling decoder with a dynamical gene regulation model.**

In this section, we present an ordinary differential equation (ODE) model which realizes the phenomenological scaling decoder for the *Drosophila* case.

The idea is fully illustrated by the toy model in Fig. 6a-c, although the gap gene network has more degrees of freedom and more scaling boundaries, and does not necessarily have to reach a dynamical attracting point. As we show below, after being properly fitted, the ODE gap gene regulation model turned out to have basically the same input-output relations as the previous phenomenological decoder. It also has satisfactory predictions on mutant patterns.

Under the local-readout hypothesis, diffusion of gap gene products is ignored. At each spatial point, the ODE model calculates the synthesis rates out of the current gap gene product ($\boldsymbol{P}$) and morphogen ($\boldsymbol{M}$) levels. (Degradation coefficient is simply set to a fixed value $\gamma$=0.05 min$^{-1}$.)

$$\frac{d}{dt}\boldsymbol{g} = \boldsymbol{f}(\boldsymbol{P}, \boldsymbol{M}) - \gamma \boldsymbol{g}$$

Here we model the regulation logics of anterior and posterior *hb* domains separately, since they are known to be generated by different regulation rules even at a course grained level [24,25]. Similar situation applies to the two *gt* domains [26,27]. Therefore, a total of 7 “gap genes” are considered.

$$\boldsymbol{g} = (g_1, \ldots, g_7) = (hb_a, hb_p, Kr, kni, gt_a, gt_p, tll)$$

Since $hb_a$ and $hb_p$ encode the same Hb protein, $\boldsymbol{P}$ has only 5 components.

$$\boldsymbol{P} = (g_1+g_2, g_3, g_4, g_5+g_6, g_7)$$

And in the input term $\boldsymbol{M}$, we consider Bcd, Tor and Caudal (Cad, which is shaped by translational repression from Bcd directly, and its profile is fitted to the measurements of Ref.[13,28]. $\mathrm{Cad} = 1/(1 + 1250 * \mathrm{Bcd}^{2.47})$).

The mHb profile is used as the initial condition of $hb_a$, and the rest $\boldsymbol{g}$ dimensions are initialized at zero.

The $\boldsymbol{f}$ term implements the known gap gene interactions reviewed in Ref.[24] (Fig. 6d and Table S2) with a set of Hill-function-like formulas defined as follows. For example, if $P_j$ plays an activating role in $f_i$, the corresponding term is:

$$f_i^j = v_{ij} \cdot \sigma\left(P_j, K_{ij}, b_{ij}\right).$$

Instead, if the regulation role is inhibitory,

$$f_i^j = 1 - \sigma\left(P_j, K_{ij}, b_{ij}\right).$$

Here, $\sigma$ is an S-shaped function with $K$ and $b$ as its parameters.

$$\sigma(x, K, b) = \frac{s\left(K(x-b)\right) - s(-Kb)}{1 - s(-Kb)}, \quad \text{where } s(z) = \frac{1}{1 + exp(-z)}$$

This function mimics the Hill function, with $K$ and $b$ together defines the steepness and half-maxima position of the S-shaped curve. These are free parameters to be fitted. Our reason for choosing this function form is purely technical – it performs better in parameters optimization by gradient descent than the original Hill function. Taken together, when a gap gene is regulated by multiple factors, the activation terms are summed and multiplied by the repression terms.

$$f_i = \left(\sum\nolimits_{Act.} f_i^j\right)\left(\prod\nolimits_{Inh.} f_i^j\right) \quad i = 1 \cdots 6$$

An exception is *tailless* (*tll*). As *tll* is known to act upstream of the gap gene network, and is shaped mainly by Tor and Bcd [29], here we explicitly write down the equation for *tll* using Hill functions.

$$f_7 = \frac{1}{1 + \left(\frac{Bcd}{0.011}\right)^4}\left(1 - \frac{1}{1 + \left(\frac{Tor}{0.07}\right)^2}\right)$$

With the above dynamical equations and initial condition, the model is integrated numerically at each embryonic position (101 discrete points along the A-P axis) independently, as we ignore diffusion of the gap gene products. The model trajectory is compared with the measured WT profiles [30] in the following two aspects to define the Loss function for parameter fitting:

(1) Averaged temporal trajectory. Those protein profiles in [30] are measured in the 14th nucleus cleavage cycle (n.c.14) at the blastoderm stage. We smoothed those profiles both spatially and temporally, and extracted 7 equally-spaced frames between 8 and 41 minutes into n.c.14. (The $t$=41min frame is the one shown in Fig. 2b of main text). The model trajectory for a $L$=1 embryo is compared with these frames at the corresponding time steps, defining the first term in Loss function.

(2) Scaling. We simulate a batch of 16 embryos with $L$'s sampled from the normal distribution $N(0, 0.15)$. Obviously, these embryos may have different gap gene patterns in the initial simulation steps due to their different initial profiles of *hb*. Therefore, to impose scaling, we add a second Loss term penalizing pattern differences *only* at the final frame ($t$=41 min into n.c.14).

Parameter fitting is then carried out using the Adam optimizer implemented in Tensorflow. Note that no mutant information is used for fitting. Table S2 lists a typical fitted parameter set.

To evaluate the fitted model, we generate a set of WT morphogen profiles with $L$ ranging from 0.85 to 1.15 according to Eqn. 5 (Fig. S11a). The ODE model successfully evolves to yield final patterns that scale with embryo lengths (Fig. S11b), and it indeed shows the overall structure quite similar to the phenomenological decoder (Fig.6e, f, main text). This is expected, as we have shown previously that any scaling local decoding scheme should follow the structure of our phenomenological decoder. Fig.6e shows the Tor=0 section (corresponding to the α plane of Fig.3a). Here, colors stand for the high-expressing gap gene given by the gene-circuit dynamics at the "final" timepoint. Dashed black lines represents the same linear classification planes as in Fig. 3a. Also note that in Fig. 6e-f, outside the WT region, domain boundaries of this ODE model indeed extend in an almost linear manner, supporting our linear extrapolation hypothesis.

As expected, having the correct decoder geometry for scaling should naturally lead to correct predictions on mutants (Fig. S11c, even gap gene mutants (Fig. S11e) which is not covered by the phenomenological decoder framework). To further evaluate these predictions in a quantitative way, we compared the predicted and measured positions of each gap gene expression domain (peaks and boundaries) in 9 different maternal morphogen mutants (*bcd*$^-$, *nos*$^-$, *tor*$^-$, *bcd1X*, *bcd4X*, *bcd*$^-$*nos*$^-$, *bcd*$^-$*tor*$^-$, *nos*$^-$*tor*$^-$, *mhb*$^-$*tor*$^-$), and the root mean square error is only 3.3% embryo length (EL). This is an impressive result. To our knowledge, it has not been reported in previous literatures that a gap gene regulation model (if fitted with WT data only) can make correct predictions in these knockout mutants.

As a negative control (Fig. S11g) to emphasize the importance of the quantitative constraints imposed by scaling, we also fit this model with the scaling requirement is removed from fitting. The resulting model shows no scaling property and hence incorrect mutant predictions (Fig. S11 g, h, i)

We also implement the scaling decoder using another widely used gene circuit modeling scheme [31-33]. The only difference is in the form of the $\boldsymbol{f}$ term (with $\gamma = 0.035$ min$^{-1}$), and the regulation network itself emerged through data fitting (as the sign of the regulation matrixes $\boldsymbol{W}$ and $\boldsymbol{V}$).

$$\frac{d}{dt}g_i = \sigma\left(\sum_j W_{ij}P_j + \sum_k V_{ik}M_k + b_i\right) - \gamma g_i \;; \qquad \sigma(x) = \frac{1}{1+e^{-x+3}}$$

The resulting effective decoder structure and predictions on mutants are also similarly satisfactory (Fig. S12). See Table S3 for model parameters.

The last thing needs to be noted here is diffusion of the gap gene products, through which decoders in adjacent nuclei may communicate with each other, going beyond the “local-decoding paradigm”. However, remind that “local-decoding” is in fact a more stringent requirement. Although diffusion is a physical effect and can never be shut down in reality, it has been pointed out by the authors of “gene circuit model” [32] that their model does not actually rely on diffusion of the gap proteins – shutting down diffusion *in silico* does not affect any dynamical process of their model albeit making the resulting profile less smooth. Similarly, our ODE model here does not include the effect of diffusion, but adding a diffusion term,

$$\frac{d}{dt}\boldsymbol{g}(x) = D\nabla^2\boldsymbol{P} + \boldsymbol{f}\big(\boldsymbol{P}(x), \boldsymbol{M}(x)\big) - \gamma\boldsymbol{g}(x)\,,$$

appears only to smooth the established spatial pattern, not affecting scaling (Fig. S11k). Here, diffusion constant is set to 0.3 $\mu m^2/s$, twice as that estimated using the exponential tail of the measured gap gene protein pattern of the *FlyEX* database [12,13] (Fig. S11i, j). In Fig. S11k, the parameter sets listed in Table S2 was used, which is fitted in the absence of diffusion. The model would be compatible to even stronger diffusion if such diffusion effect is considered during parameter fitting.

**S12. Nos/mHb as the posterior gradient.**

Our work is largely inspired by the bi-gradient model. The original bi-gradient models [20,34], however, are not widely accepted partly because they introduced a hypothetical (and non-exist) posterior gradient. This hypothetical gradient is needed because Nos was ruled out from the very beginning, based on the observations by (and only by) Ref.[4] that positional variability of Hb boundary do not seem to be seriously affected in *nos*$^{-}$ or *mhb*$^{-}$ embryos. By contrast, in our model mHb/Nos *is* the second gradient. Below, we demonstrate that the observations in [4] can in fact be well explained with the updated understanding of Bcd gradient since 2002, namely, it has very low noise and its amplitude partially scales with the embryo length. Therefore, those observations cannot rule out mHb/Nos; a hypothetical posterior gradient is not actually needed.

There are two arguments relevant to this topic in the 2002 paper by Houchmandzadeh et al. [4]. First, if Nos/mHb is the factor that helps setting Hb-Kr boundary (Tor is effectively zero here) together with Bcd, then in *nos*$^{-}$ or *mhb*$^{-}$ embryos, positional noise of the Hb boundary should follow that of Bcd. The observed positional error of Hb boundary increases from 1%EL (WT) to 1.6%EL (*nos*$^{-}$), while the positional noise of Bcd measured by [4] is almost 30%EL. There seem to be a big gap. However, as has been correctly pointed out later by Gregor et al., [18], this 30% positional error of Bcd profile is an artifact due to inappropriate normalization; and the true Bcd positional error should be around 1~2% EL if measured with GFP-tagged Bcd and normalized properly [18,35] (and Fig. S2). Therefore, this first argument is invalid from the current point of view. Hb boundary noise *is* at the same level of Bcd in *nos*$^{-}$ or *mhb*$^{-}$ background.

A second related observation of Ref.[4] is that, scaling of the Hb boundary seems not to be completely destroyed in *nos*$^{-}$ or *mhb*$^{-}$ embryos: its absolute position (measured from the anterior pole, in μm, denoted as $x_{hb}$) remains to depend on embryo length (with a linear correlation coefficient about 0.7). We think this evidence is also not sufficient for excluding mHb/Nos. This correlation can be explained by the Bcd amplitude effect – that larger embryo tends to have higher absolute level of Bcd [5] (and SI-1). This amplitude effect (which is even slightly underestimated if setting $\beta$=2 as in Eqn. 5a) makes the *hb* boundary appear to be partially scaling, even if it is determined solely by Bcd threshold:

$$\left.\frac{\partial x_{hb}}{\partial L}\right|_{L=1} = \beta\lambda_B \approx 0.33 \qquad (S12.1)$$

The *hb* boundary locates near the center of embryo. Thereby this quantity should be approximately 0.5 for perfect scaling, and equals to 0 for non-scaling as naively expected. The seemly partially preserved scaling in *nos*$^-$ or *mhb*$^-$ can be explained in this way.

**S13. Discussion on the long-germband insect *Megaselia abdita*.**

Geometrically speaking, in our framework scaling stems from quantitative matching of the decoder decision boundaries with the *y*-constant curves. On the other hand, from the regulatory perspective, it is equivalent to say that all three morphogen levels change as *L* varies, but their effects on the entire gap gene network should cancel out (at the linear order) to give unchanging outputs. This kind of precise cancellation relies on quantitative tuning of regulatory link strengths in the gap gene network. Such "fine-tuning" may not seem to be a reliable mechanism at a first glance. However, it is consistent with the current understanding of the *evolution* of the gap gene network. The maternal morphogen system is very diverse among different long germband insects, but gap gene cross regulation network is much more conserved [24]. We suggest that by tuning its link strengths quantitatively, a long-germband insect species can easily make the "ancient" gap gene network adapt to its specific maternal morphogen system and achieve scaling patterning.

We would like to briefly discuss, with our scaling framework, another long-germband insects with different sets of maternal morphogen gradients – *Megaselia abdita*. As experimental data (especially on mutants) are very limited, we cannot discuss this point in depth. Like *Drosophila*, *Megaselia* also have Bcd and mHb. The mHb profile is very similar to that of *Drosophila*, but Bcd extends more to the posterior [36,37]. Different morphogen shapes lead to different geometry of the WT point clouds (Fig. S13a), hence different predictions on mutants. In the case of more extended Bcd, when the decision boundary is linearly extrapolated to predict *bcd*$^{-}$, the *kni* domain expands and the *Kr* domain disappears completely (Fig. S13 illustrates this point with $\lambda_{\mathrm{Bcd}}$=0.225), which is qualitatively the situation observed in *bcd*$^{-}$ *Megaselia* [38].

**Supplemental Figure legends.**

**Fig. S1.** The mHb profile can be fitted by a sigmoidal curve. The Hb protein profile in n.c.12 embryos are regarded as maternal Hb here. Data cited from the *FlyEX* database.

**Fig. S2.** Simulating noise in the Bcd gradient by a Poisson noise term. Black curve: positional error (standard deviation $\sigma_y$) calculated according to Eqn. S2.2 (with $N_0$=1000, $\lambda_B$=0.165, and $\beta$=2). Blue/green data points: the measured Bcd positional error [18], without/with the known measurement error being subtracted. The Poisson noise term simulates Bcd noise correctly in the region $0.2<y<0.6$.

**Fig. S3.** Predicting mutant fate-map with linear classifiers. (**a**) A graphical illustration of predicting the fate of $y_{\mathrm{mut}}$=0.4 point in *bcd*$^-$*tor*$^-$ mutant. Grey and white pixels stand for different classifier outputs on this point. (**b**) A typical case where the "posterior dominant rule" should be employed. A linear classification plane fitted at the anterior (#20 here) intersected with much more posterior parts of the WT point cloud, far away from where the #20 plane was fitted. Hence its classification on those posterior points (yellow rectangle) should be ignored. (**c**) By listing the outputs of all classification planes on all points form a mutant embryo, the predicted fate map is the grey-white boundary. Those outputs ignored by the "posterior dominant rule" are shown in lighter color.

**Fig. S4.** Effects of the Bcd $\beta$ factor and noise amplitude on model performance. (**a**) By re-adjusting the classification orientations, a scaling phenomenological decoder is also obtained without the Bcd amplitude factor. It finds the correct $\tilde{y}$ values with relatively small error (RMSE~1%) for WT embryos with length variations. (**b**-**c**) Though scaling is not affected by dropping the amplitude factor, the geometry of the

WT point cloud hence the corresponding mutant phenotypes do change (marked by the arrowhead in panel **c**). (**d**) Setting $\beta$ to 3 does not have much influence on the mutant predictions, compared with Fig. 3d of the main text. (**e-f**) With the settings in the main text ($\beta$=2), changing the Poisson noise strength do not affect our main results. The predictions are still satisfactorily consistent with experiments.

**Fig. S5.** The Bayesian decoder. (**a**) Decoding error for the WT ensemble in Fig.2d of main text. (**b**) The Root-Mean-Squared decoding error is on the same order as our linear decoder. Though arbitrary nonlinear boundary geometry is allowed by the Bayesian decoder, its error is even larger than the linear one. (**c**) Bayesian decoder outputs. Within the region covered by the WT point cloud, they are quite similar to the linear decoder outputs. While there is much difference away from the WT point cloud. Dashed black lines are the same as Fig.3a in the main text. (**d**) Tangential views of the iv-v and v-vi *linear* classification planes of the decoder used in main text. The remaining classification error is due to the Poisson noise rather than nonlinearity in the domain interface geometry.

**Fig. S6.** Outputs of the scaling decoder on mHb=0 and mHb=$mHb_0$ planes. The 3-d orientations of the decision planes naturally explans that while *nos*$^-$ embryos have lost the abdominal fates (domains v & vi), *nos*$^-$*mhb*$^-$ embryos still have these two domains just as WT.

**Fig. S7.** Maternal morphogen mutant embryos with greatly changed length. (**a**) A graphical illustration of the prediction in Fig. 4a of the main text. (**b**) Measured gap gene domain positions in normal-length and greatly shortened *bcd*$^-$ embryos[7]. The narrow anterior *gt* domain is not shown here. (**c**) Overlapping panels A and B for comparison. (**d**) Predicted and measured gap gene domain boundaries in greatly shortened *bcd6X* embryos. Solid lines, model prediction; dots, measurements by Ref.[7].

(**e**) Analytical calculation of the *hb* boundary position in *bcd6X* embryos of varying lengths. The black curve follows Eqn. S7.2, with parameters $\lambda_B$=0.165, and $\beta$=2. (**f**) Predicted gap gene domain positions of many other maternal morphogen mutants with greatly changed $L$'s. The horizontal dashed lines mark typical range of natural length variation within a fly line.

**Fig. S8.** The three maternal gradients function as two bi-gradient pairs. (**a-c**) Size sensitivity $S_L$ evaluated at $L$=1. This quantity is equivalent to absolute slopes of the domain boundaries in Fig. 5a, c, d in the main text. $S_L$ is low everywhere in the WT embryo, indicating good scaling (black, $S_L$ <0.1). Missing Bcd (**a**) destroys scaling completely (purple), while missing the posterior (**b**) or terminal (**c**) morphogen only affects scaling in part of the embryo. The dashed grey lines show the non-scaling baselines of $S_L$. (**d**) Contribution of each morphogen (defined in Section S8) in discriminating adjacent positions in WT embryos. Bcd forms bi-gradient pair with mHb in the middle part, and with Tor near both ends. Note that morphogen contributions are different for different embryo lengths. Nots that mHb seems to play a more important role in larger embryos. (**e**) Therefore, the same perturbation in Nos should introduce more sever segmentation defects for larger embryos. Here, we present predictions on larger and smaller embryos with reduced Nos dosage/activity. Gap gene pattern should be basically normal-looking for $L$=0.7, while for $L$=1.3 the *Kr* domain should expand greatly. This prediction may be checked experimentally in the future.

**Fig. S9.** Shift of the *even-skipped* (*eve*) stripes under Bcd dosage change. (**a**) Predictions with or without the other two maternal morphogens. (**b**) A semi-quantitative comparison with the experimental measurements in Ref.[22].

**Fig. S10.** Optimal decoder for noisy gradients with or without length variation. Here, “scaling decoder” stands for the model studied in the main text. The “noise-only decoder” is defined similarly – *L* is fixed to 1 in generating the WT point cloud, while the Poisson noise term is the same as the scaling decoder. “Null model” is the most naïve form of linear decoder, that the decision planes are always defined to be normal to the (Bcd, mHb, Tor) curve. (**a-h**) Geometry, contribution of each morphogen, and performance in decoding the full-version WT point cloud (i.e., that in Fig.2d main text) of the noise-only decoder and the null model They differs from the scaling decoder (**i-l**) significantly. (**m-o**) The scaling decoder turns out to be a better description of the real *Drosophila* gap gene system; its predictions (grey-white boundary, as in Fig. S3) on mutant patterns match the best with experiments (black dotts). For the noise-only decoder and the null model, where their prediction deviates from the experiments are marked by red arrowheads. (**p-r**) According to the scaling decoder, Bcd should play a non-negligible role even near the posterior pole. This is supported by experiments (expansion of the posterior *hb* domain in *bcd*$^-$). Note that this feature is not captured by either the null model or the noise-only decoder. (**s-v**) Analysis of the noise-only decoder and the null model following Fig. 5a and f-h of the main text.

**Fig. S11.** The ODE-based gap gene regulation model. (**a**) The morphogen profiles for *L* ranging from 0.85 to 1.15 following Eqn. 5. These morphogens define the external inputs and initial condition of an ordinary differential equation model for the gap gene network. Note that mHb is multiplied by a factor 4 for better visualization. (**b**) This gene regulation model generates scaling gap gene pattern by reading the non-scaling morphogens in panel A. The requirement of scaling in included in parameter fitting. (**c**) Mutant Predictions (solid lines) and corresponding measured profiles (dashed lines, cited from Refs.[6] and[21]). (**d**) A systematic assessment of mutant predictions. Predicted vs. measured positions of peaks and boundaries of the gap gene domains are shown. The predicted positions have a root-mean-square-error (RMSE) of around 3.3% embryo length, which is quite impressive. (**e**) This model even has reasonable predictions on gap gene mutants. Take the $Kr^-$ mutant as an example here. Semi-quantitatively, without *Kr*, the posterior *gt* domain should expand anteriorly and eliminate the *kni* domain by inhibition. This is exactly the situation observed in experiments [26,39]. (**f-h**) Regulation network topology alone cannot ensure scaling. (**f**) The ODE model failed to achieve scaling if scaling is not explicitly introduced in parameter fitting. (**g-h**) When being viewed as a decoder of maternal morphogens, its structure deviates from that of our phenomenological scaling decoder (dashed lines). Having the "correct" regulation network topology (identical to Fig. 6d) is not enough for scaling. Quantitative features matter. (**i-j**) Estimating diffusion constant for the gap gene products using the "exponential tails" in the protein profiles. Half-life of the Hb or Kr protein is assumed to be 14 min. Note that this is only a rough estimation. (**k**) The scaling ODE model of panel B is robust to diffusion. The resulting pattern remains normal and scaling, albeit smoothed by diffusion. Diffusion constant here is relatively large, twice the estimated value.

**Fig. S12.** The scaling gene circuit model. (**a**) With its parameters being properly fitted, using the Loss function defined in Section S11, the gene circuit model generates scaling output pattern successfully. (**b**) The equivalent decoder structure also follows the phenomenological linear decoder (dashed lines). (**c**) The gap gene cross regulation network emerges from data fitting. (**d-e**) This gene circuit model also has satisfactory predictions on maternal morphogen mutants, as expected. (**f**) It can also reproduce Fig. 5a-d of the main text, reflecting its structural similarity with the phenomenological scaling decoder.

**Fig. S13.** Generating scaling *Drosophila* gap gene pattern with a more extended Bcd profile. (**a**) Increasing the Bcd length constant (form 0.165 used in main text to 0.225

here) leads to "stretching" of the WT point cloud. The decision boundaries of a scaling decoder are also shifted (dashed black lines). (**b**) As a result, by extrapolating with these decision planes, the fate corresponding to the *Kr* expression domain (iv) no longer presents on the Bcd=0 plane. Compare this panel with Fig. 3a-b to see this difference. Therefore, in a semi-quantitative sense, $bcd^-$ embryo should no longer have a *Kr* domain with this more extended Bcd. Note that the *Drosophila* gap gene positions are used in this figure, making it comparable with Figs. 2 and 3 in the main text.

**Table S1.** Peak and boundary positions extracted form maternal morphogen mutants.

| *wt* | | *bcd1X* | | *bcd4X* | | $bcd^{-}$ | | $nos^{-}$ | |
|---|---|---|---|---|---|---|---|---|---|
| Pos (% EL) | name | pos | name | pos | name | pos | name | pos | name |
| 12.8 | hb1L | 14.8 | hb1L | 15.6 | hb1L | 18.1 | hb2P | 18.2 | hb1L |
| 47.4 | hb1R | 40.8 | hb1R | 56.6 | hb1R | 11.5 | hb2R | 50.5 | hb1R |
| 80.8 | hb2P | 80.7 | hb2P | 84.8 | hb2P | 25.8 | hb2L | 82.8 | hb2P |
| 74.8 | hb2L | 74.2 | hb2L | 79.4 | hb2L | 77.7 | hb2P | 74.6 | hb2L |
| 87.7 | hb2R | 86.5 | hb2R | 89.3 | hb2R | 68 | hb2L | 89 | hb2R |
| 51.6 | krP | 46.5 | krP | 60.6 | krP | 86.2 | hb2R | 56.6 | krP |
| 45.8 | krL | 40.3 | krL | 55.4 | krL | 39.4 | krP | 50 | krL |
| 58.5 | krR | 54.3 | krR | 66.1 | krR | 32.2 | krR | 67.4 | krP |
| 61.6 | kniP | 58.5 | kniP | 68.7 | kniP | 47.1 | krR | 38.3 | gt1P |
| 56.5 | kniL | 52.5 | kniL | 64.4 | kniL | 47.4 | kniP | 23.8 | gt1L |
| 66.6 | kniR | 63.9 | kniR | 72.7 | kniR | 35.8 | kniL | 43.5 | gt1R |
| 35.4 | gt1P | 28.3 | gt1P | 44.4 | gt1P | 55.6 | kniR | | |
| 20 | gt1L | 18.9 | gt1L | 26.6 | gt1L | 26.3 | gt2P | | |
| 40.9 | gt1R | 33.5 | gt1R | 49.4 | gt1R | 21.9 | gt2R | | |
| 69.7 | gt2P | 67.7 | gt2P | 74.8 | gt2P | 31.5 | gt2L | | |
| 64.7 | gt2L | 62.6 | gt2L | 71 | gt2L | 58.6 | gt2P | | |
| 74.8 | gt2R | 73.5 | gt2R | 79.9 | gt2R | 50.8 | gt2L | | |
| 8 | otdL | | | | | 69 | gt2R | | |
| 26 | otdR | | | | | | | | |
| 22 | emsL | | | | | | | | |
| 31 | emsR | | | | | | | | |
| 25 | btdL | | | | | | | | |
| 33 | btdR | | | | | | | | |

Table S1 continued.

| *tor-* | | *bcd-tor-* | | *nos-tor-* | | *bcd-nos-* | | *nos-tor-mhb-* | |
|---|---|---|---|---|---|---|---|---|---|
| pos | name | pos | name | pos | name | pos | name | pos | name |
| 48.4 | hb1R | 10 | krP | 3.32 | hb1L | 17.2 | hb2P | 42.7 | hb1R |
| 53.6 | krP | 29 | krP | 49.6 | hb1R | 11.2 | hb2R | 46.7 | krP |
| 47.5 | krL | 47.1 | krR | 60 | krP | 25.3 | hb2L | 40 | krL |
| 60.9 | krR | 51.6 | kniP | 80 | krP | 82.8 | hb2P | 55 | krR |
| 65.6 | kniP | 43.1 | kniL | 49.3 | krL | 74.8 | hb2L | 60 | kniP |
| 59.2 | kniL | 62.6 | kniR | 33.3 | gt1P | 89.4 | hb2R | 53 | kniL |
| 72.5 | kniR | 55.5 | gt2L | 9.9 | gt1L | 50 | krP | 69.7 | kniR |
| 35.4 | gt1P | 70 | gt2P | 39.3 | gt1R | 33.2 | krR | 30 | gt1P |
| 11.2 | gt1L | | | | | 66.6 | krR | 11.1 | gt1L |
| 41.1 | gt1R | | | | | 25.2 | gt2P | 35.4 | gt1R |
| 78.8 | gt2P | | | | | 20.2 | gt2R | 76 | gt2P |
| 70.7 | gt2L | | | | | 29.2 | gt2L | 66 | gt2L |
| | | | | | | 74.8 | gt2P | | |
| | | | | | | 70.5 | gt2L | | |
| | | | | | | 80 | gt2R | | |

| 6Bvas-exu- | |
|---|---|
| pos | name |
| 40 | otdR |
| 50 | emsR |
| 56 | btdR |
| 24 | emsL |
| 30 | btdL |
| 82 | otdR |
| 80 | emsR |
| 78 | btdR |
| 90 | emsL |
| 85 | btdL |

**Table S2.** Parameters for the scaling ODE model.

| Regulation link | Sign | W | b | c |
|---|---|---|---|---|
| Hb to hb-a | Act. | 12.86 | 8.1300 | 0.2665 |
| Kr to hb-a | Inh. | 9.674 | -3.5490 | / |
| Kni to hb-a | Inh. | 33.98 | 10.6600 | / |
| Bcd to hb-a | Act. | 27.03 | 6.7250 | 0.8884 |
| Tor to hb-a | Inh. | 3.265 | 9.8870 | / |
| Kr to hb-p | Inh. | 16.35 | 8.4220 | / |
| Kni to hb-p | Inh. | 17.12 | 8.8830 | / |
| Tll to hb-p | Act. | 32.97 | 9.9490 | 2.2230 |
| Tor to hb-p | Inh. | 8.948 | 10.9600 | / |
| Hb to Kr | Inh. | 5.909 | -0.3211 | / |
| Kni to Kr | Inh. | 11.4 | -2.7410 | / |
| Gt to Kr | Inh. | 20.2 | 2.4620 | / |
| Tll to Kr | Inh. | 25.35 | 9.1960 | / |
| Bcd to Kr | Act. | 8.417 | -0.6508 | 5.7930 |
| Cad to Kr | Act. | 20.05 | -3.6540 | 1.5560 |
| Tor to Kr | Inh. | 24.06 | 12.2500 | / |
| Hb to kni | Inh. | 38.25 | 4.3050 | / |
| Gt to kni | Inh. | 24.56 | -2.7790 | / |
| Tll to kni | Inh | 69.30 | 10.7100 | / |
| Bcd to kni | Act. | / | / | 0 |
| Cad to kni | Act. | 6.008 | -0.6122 | 2.0760 |
| Tor to kni | Inh. | 3.993 | 6.9190 | / |
| Kr to gt-a | Inh. | 22.08 | 7.3770 | / |
| Bcd to gt-a | Act. | 27.1 | 2.0770 | 1.5040 |
| Tor to gt-a | Inh. | 27.27 | 1.8270 | / |

| | | | | |
|---|---|---|---|---|
| Hb to gt-p | Inh. | 21.55 | 12.3600 | / |
| Kr to gt-p | Inh. | 14.45 | 2.8210 | / |
| Tll to gt-p | Inh. | 4.086 | 9.1700 | / |
| Cad to gt-p | Act. | 6.042 | -3.1790 | 3.3710 |
| Tor to gt-p | Inh. | 7.685 | 1.8580 | / |

**Table S3.** Parameters for the scaling gene circuit model.

Weights: W&V

| | *hb-a* | *hb-p* | *Kr* | *kni* | *gt-a* | *gt-p* |
|---|---|---|---|---|---|---|
| Hb | -0.855 | 4.031 | -3.526 | -15.49 | 1.084 | -22.06 |
| Kr | 1.179 | -22.44 | 4.152 | -2.664 | -14.11 | -18.03 |
| Kni | -64.61 | -11.24 | -3.721 | 6.641 | -37.88 | 3.473 |
| Gt | 2.135 | 4.918 | -10.06 | -9.866 | -0.609 | 2.392 |
| Tll | -17.6 | -0.248 | -14.01 | -44.98 | 1.735 | -2.996 |
| Bcd | -2.911 | -38.12 | -3.172 | -8.143 | -15.35 | -31.18 |
| Cad | -3.129 | 4.412 | -1.654 | 2.084 | -8.448 | 5.898 |
| Tor | -2.928 | -3.23 | -51.2 | -4.751 | -20.07 | -5.281 |

Bias:

| | *hb-a* | *hb-p* | *Kr* | *kni* | *gt-a* | *gt-p* |
|---|---|---|---|---|---|---|
| Bias b | 5.127 | -2.953 | 4.732 | 1.728 | 7.834 | -1.639 |
