## Supplemental Figures for "Scaling dictates the decoder structure"

### Fig. S1

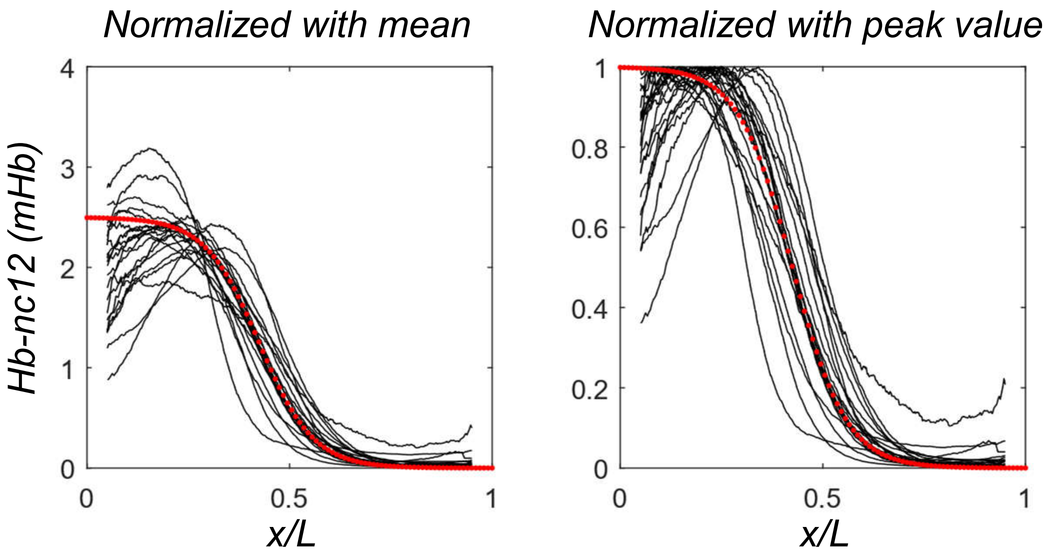

Normalized with mean
Normalized with peak value
Hb-nc12 (mHb)
4
3
2
1
0
0
0.5
1
x/L
0.8
0.6
0.4
0.2


### Fig. S2

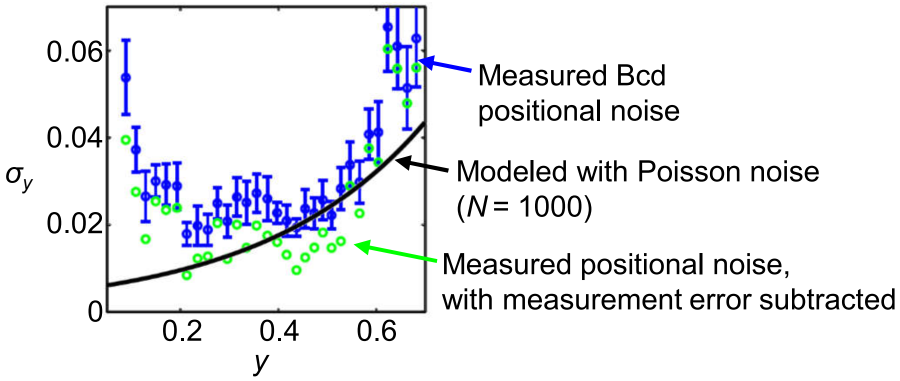

0.06
0.04
0.02
0
$\sigma_y$
0.2
0.4
0.6
y
Measured Bcd positional noise
Modeled with Poisson noise ($N$ = 1000)
Measured positional noise, with measurement error subtracted

**Fig. S3**

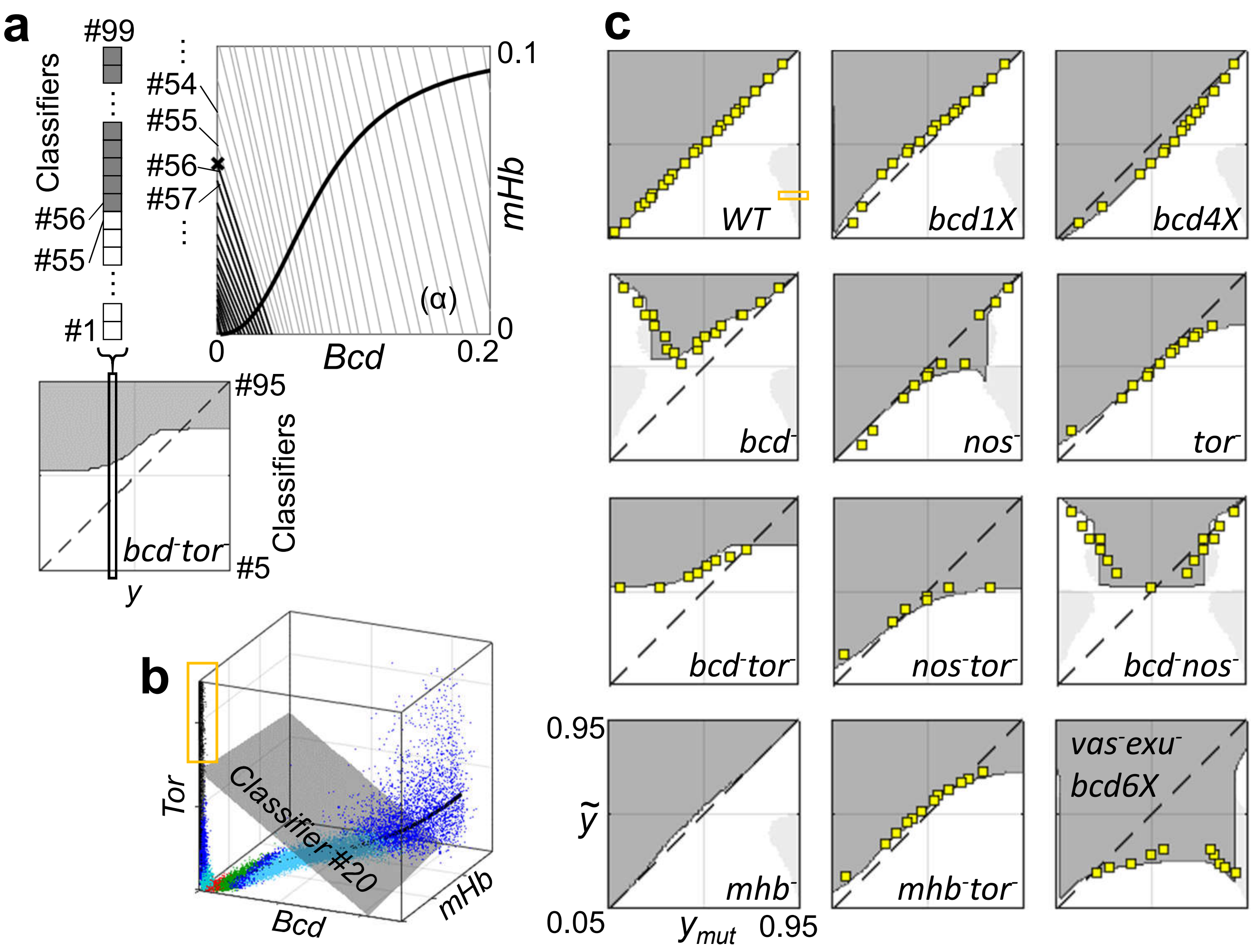
a
#99
Classifiers
#56
#55
#1
#54
#55
#56
#57
0.1
0
(α)
0
Bcd
0.2
mHb
#95
Classifiers
#5
bcd-tor-
y
b
Tor
Classifier #20
Bcd
mHb
c
WT
bcd1X
bcd4X
bcd-
nos-
tor-
bcd-tor-
nos-tor-
bcd-nos-
0.95
0.05
ỹ
mhb-
mhb-tor-
vas-exu-
bcd6X
y mut
0.95

### Fig. S4

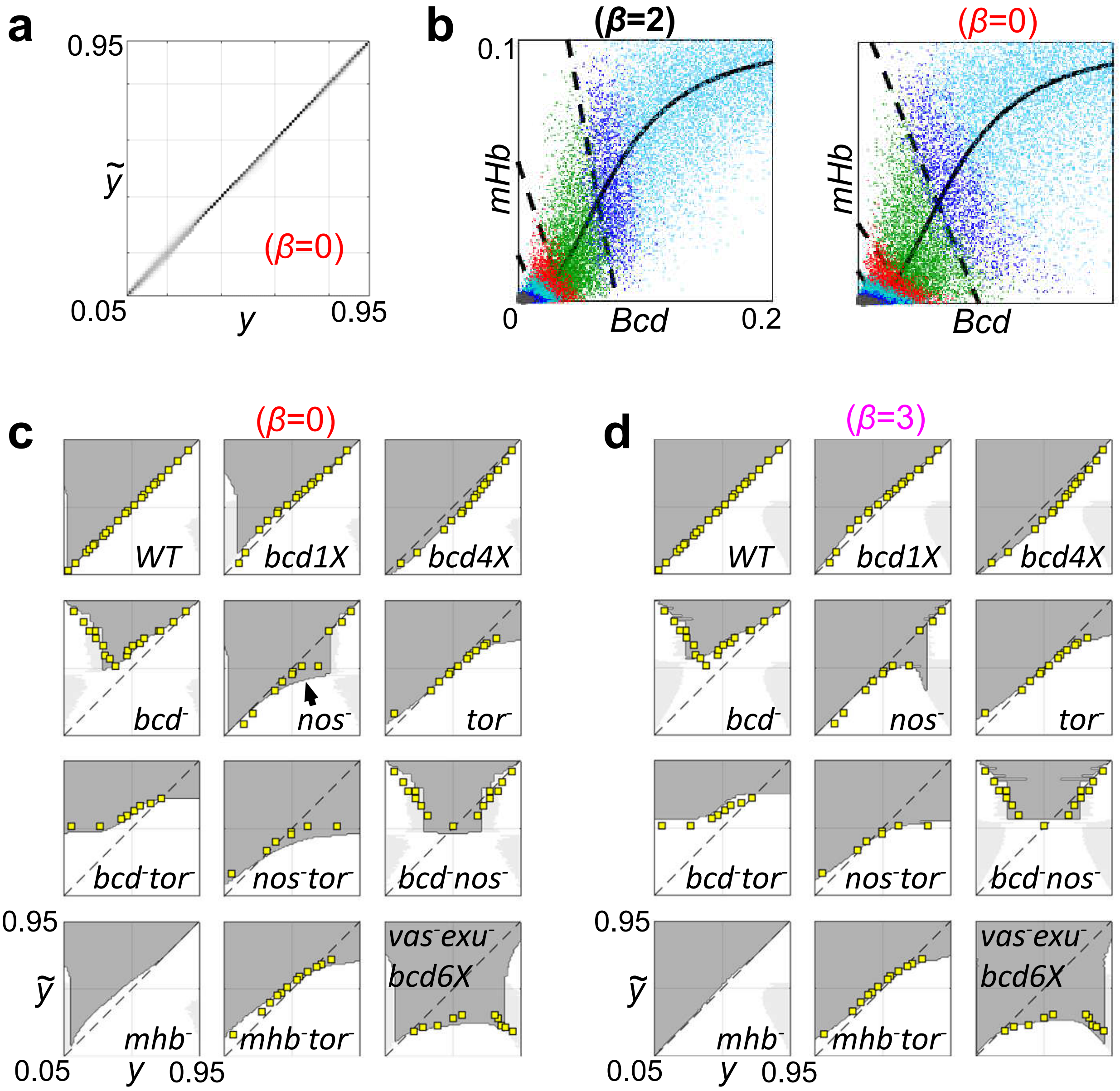
a
0.95
ỹ
(β=0)
0.05
y
0.95
b
(β=2)
0.1
mHb
0
Bcd
0.2
(β=0)
mHb
Bcd
c
(β=0)
WT
bcd1X
bcd4X
bcd-
nos-
tor-
bcd-tor-
nos-tor-
bcd-nos-
0.95
ỹ
mhb-
mhb-tor-
vas-exu-
bcd6X
0.05
y
0.95
d
(β=3)
WT
bcd1X
bcd4X
bcd-
nos-
tor-
bcd-tor-
nos-tor-
bcd-nos-
0.95
ỹ
mhb-
mhb-tor-
vas-exu-
bcd6X
0.05
y
0.95

### Fig. S4-continued

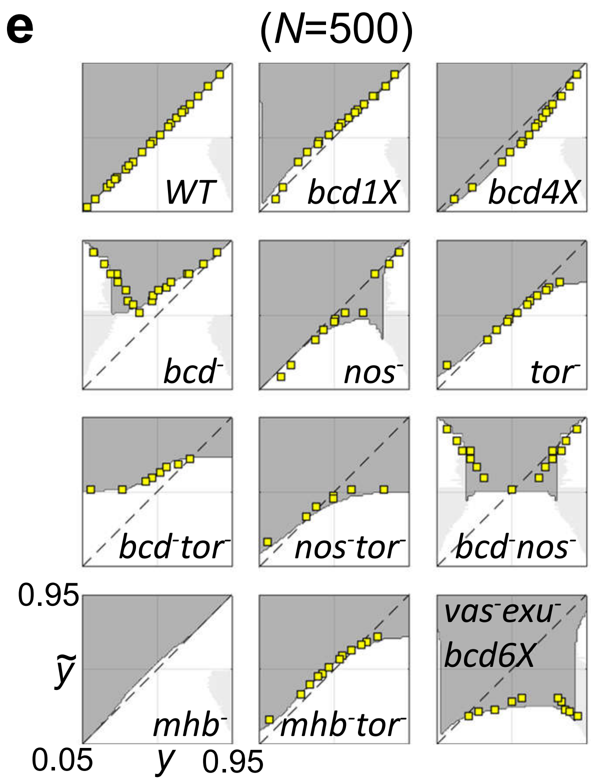


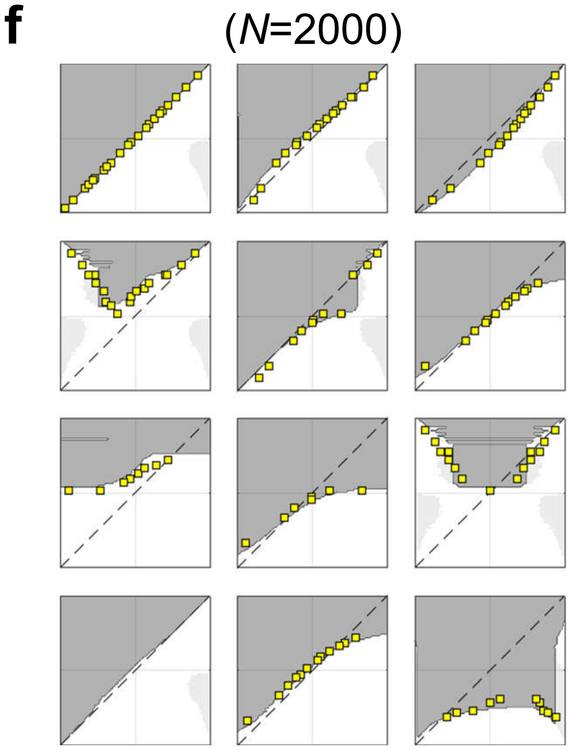

#### Fig. S5

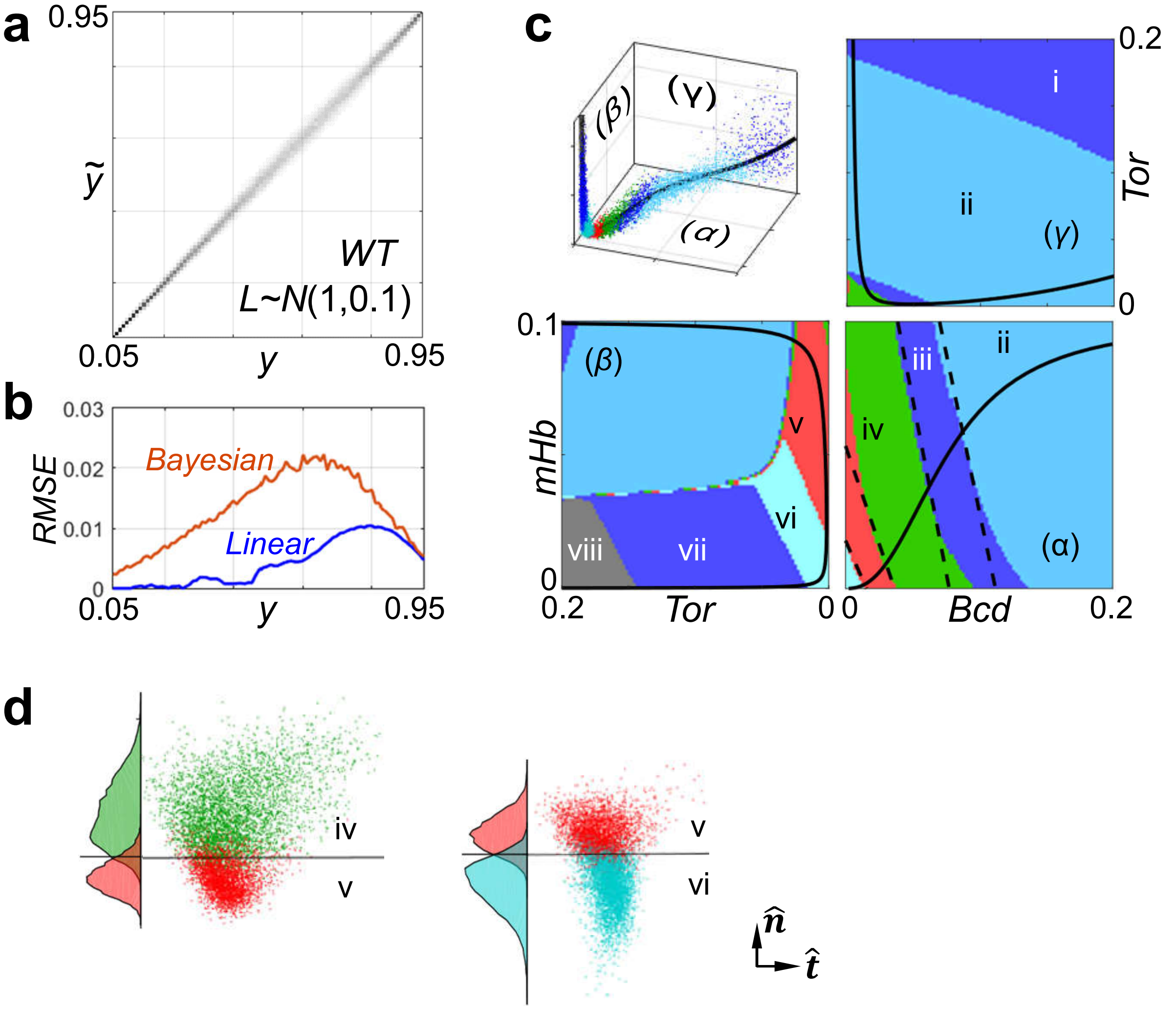

a
0.95
$\tilde{y}$
WT
L~N(1,0.1)
0.05
y
0.95
b
0.03
RMSE
0.02
0.01
0
Bayesian
Linear
0.05
y
0.95
c
(β)
(γ)
(α)
i
ii
(γ)
Tor
0.2
0
0.1
mHb
(β)
v
vi
vii
viii
0
0.2
Tor
0
0
Bcd
0.2
iii
ii
iv
(α)
d
iv
v
v
vi
$\hat{n}$
$\hat{t}$


#### Fig. S6

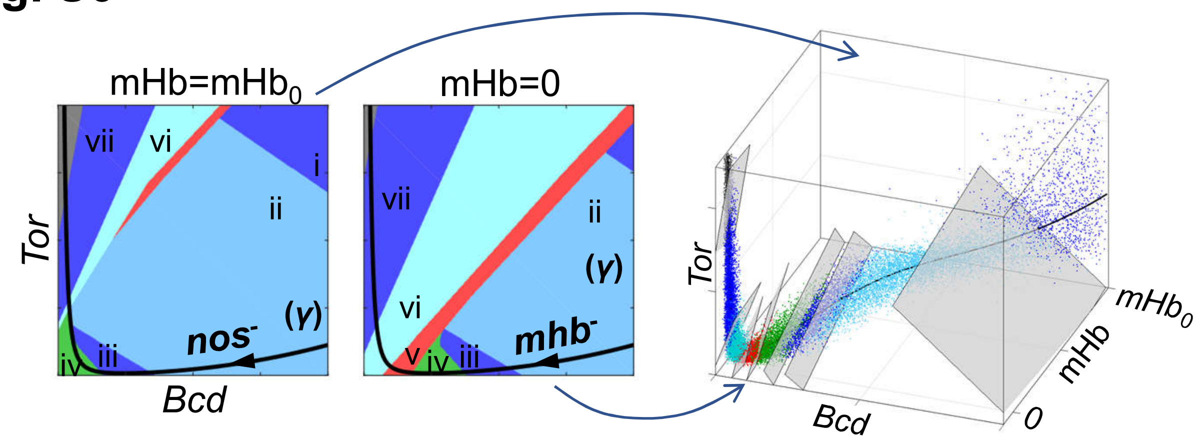

mHb=mHb0
mHb=0
vii
vi
i
ii
Tor
(γ)
nos-
iv
iii
Bcd
vii
ii
(γ)
vi
mhb-
v
iv
iii
Tor
Bcd
mHb0
mHb
0

### Fig. S7

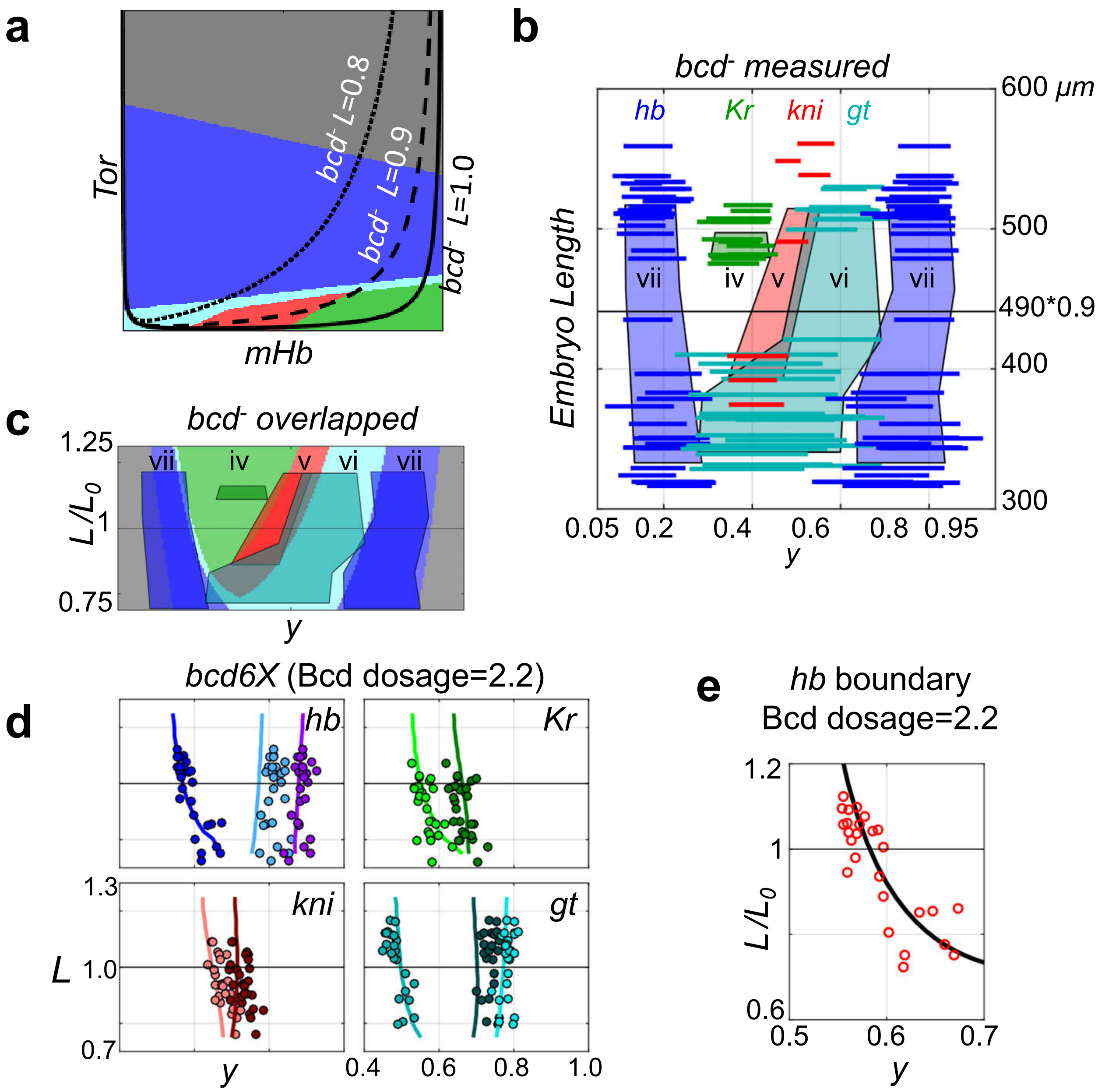

a
Tor
mHb
bcd- L=0.8
bcd- L=0.9
bcd- L=1.0
b
bcd- measured
hb
Kr
kni
gt
Embryo Length
600 µm
500
490*0.9
400
300
vii
iv
v
vi
vii
0.05 0.2 0.4 0.6 0.8 0.95
y
c
bcd- overlapped
L/L0
1.25
1
0.75
y
bcd6X (Bcd dosage=2.2)
d
hb
Kr
kni
gt
L
1.3
1.0
0.7
y
0.4 0.6 0.8 1.0
e
hb boundary
Bcd dosage=2.2
L/L0
1.2
1
0.6
0.5 0.6 0.7
y

### Fig. S7-continued

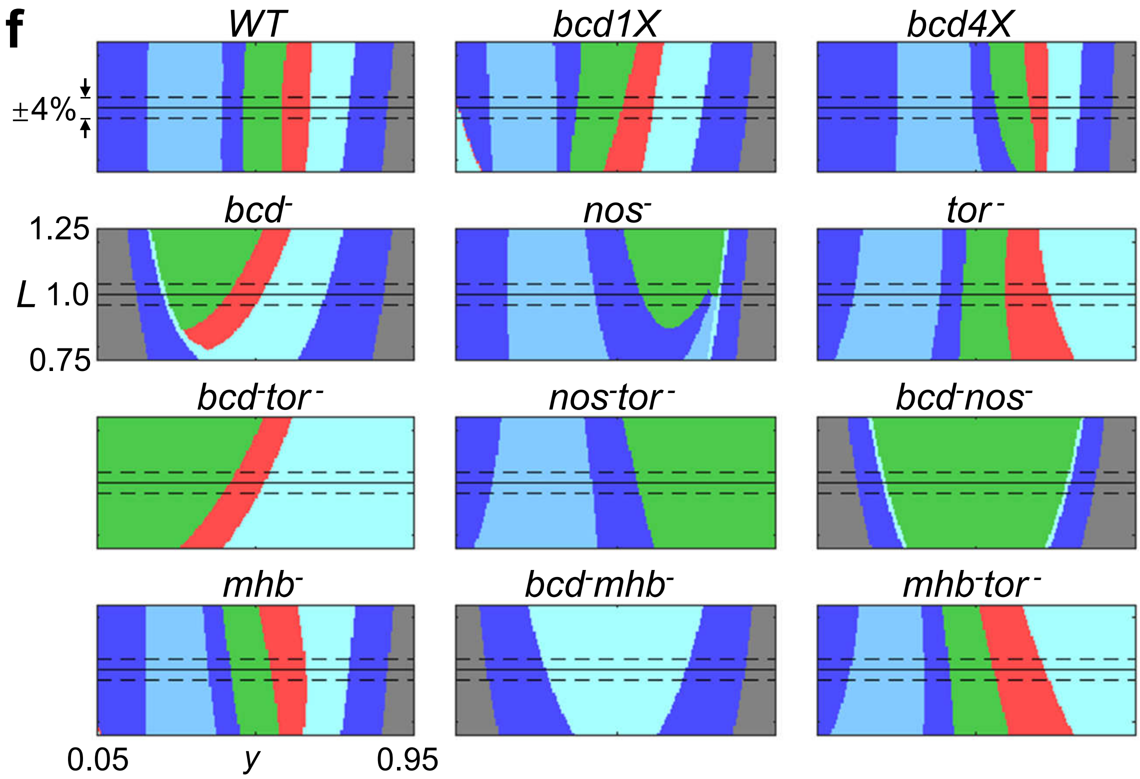

#### Fig. S8

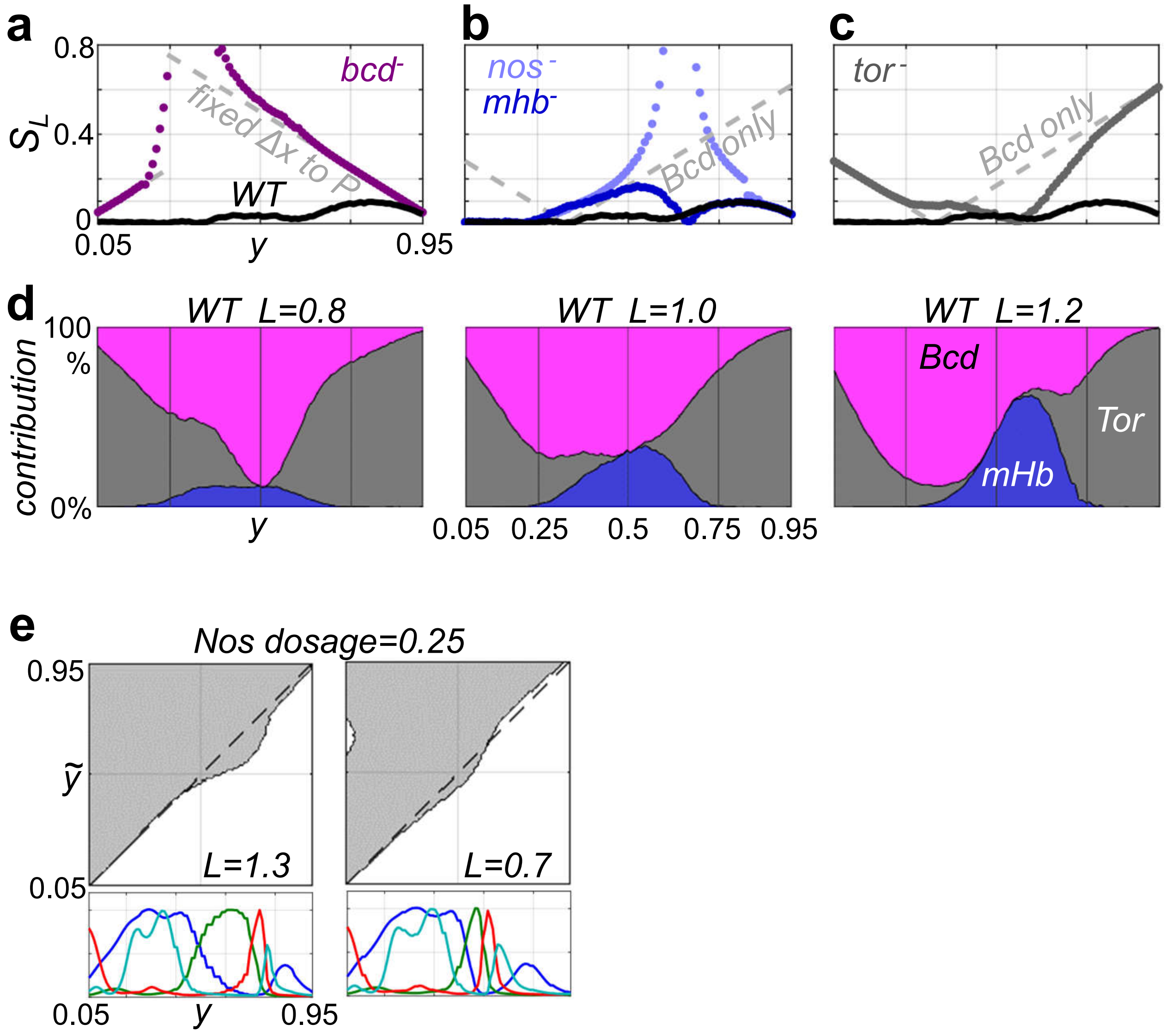

a
b
c
0.8
$S_L$ 0.4
0
bcd-
fixed Δx to P
WT
0.05
y
0.95
nos-
mhb-
Bcd only
tor-
Bcd only
d
WT L=0.8
WT L=1.0
WT L=1.2
100 %
contribution
0%
y
0.05 0.25 0.5 0.75 0.95
Bcd
Tor
mHb
e
Nos dosage=0.25
0.95
ỹ
0.05
L=1.3
L=0.7
0.05
y
0.95

#### Fig. S9

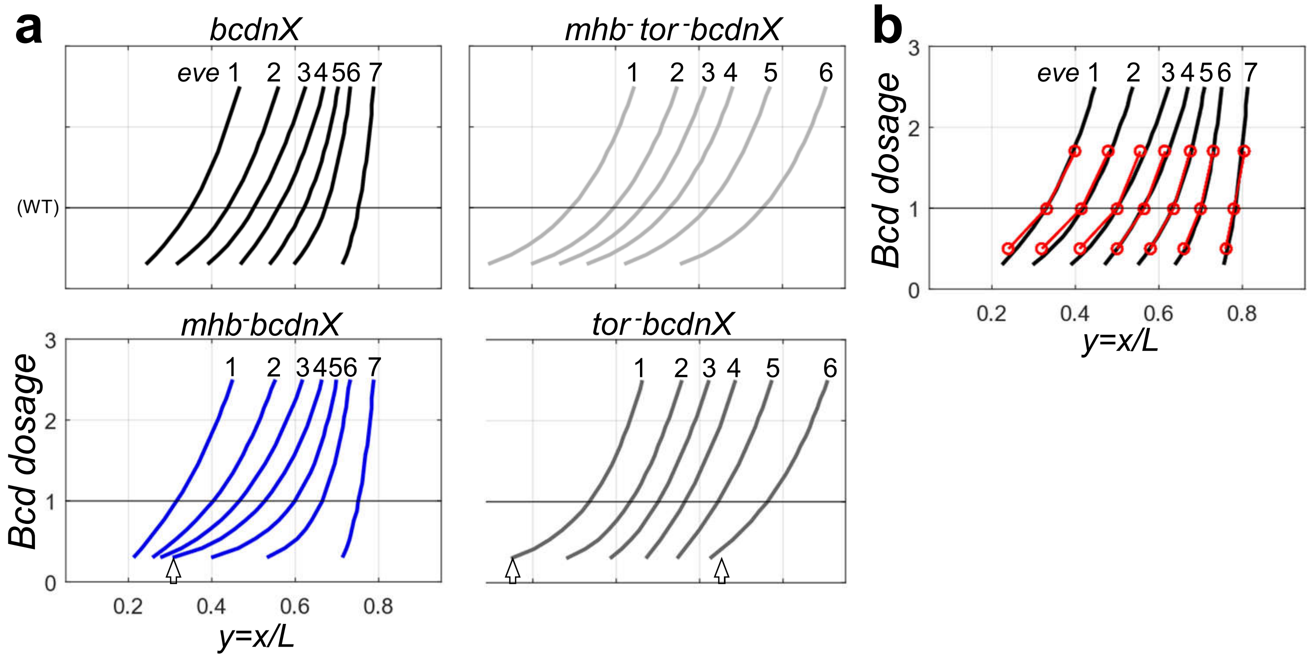

a
bcdnX
eve 1 2 3 4 5 6 7
(WT)
mhb⁻ tor⁻bcdnX
1 2 3 4 5 6
mhb⁻bcdnX
1 2 3 4 5 6 7
tor⁻bcdnX
1 2 3 4 5 6
Bcd dosage
0 1 2 3
0.2 0.4 0.6 0.8
y=x/L
b
eve 1 2 3 4 5 6 7
Bcd dosage
0 1 2 3
0.2 0.4 0.6 0.8
y=x/L

#### Fig. S10

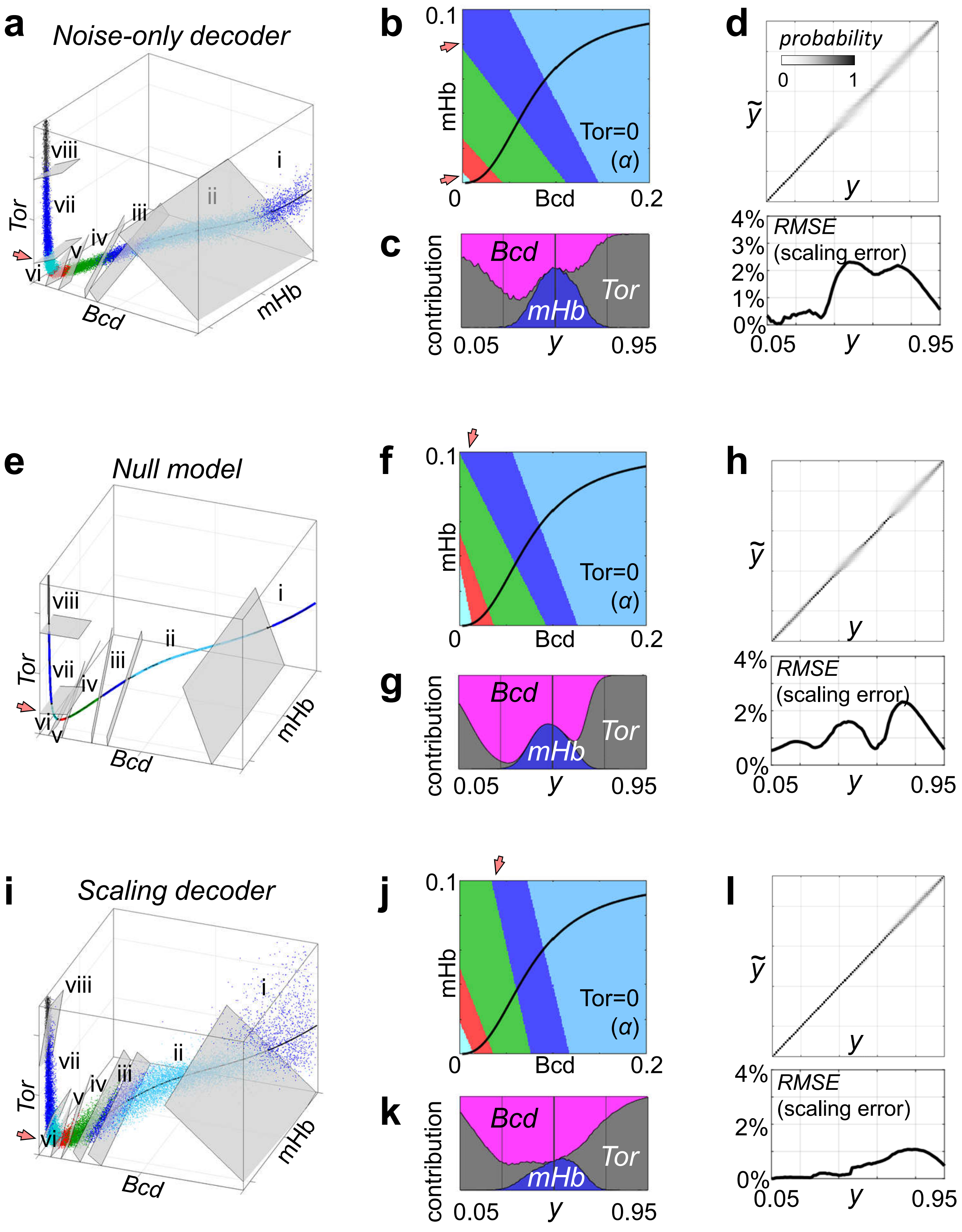
a
Noise-only decoder
viii
vii
i
ii
iii
iv
v
vi
Tor
Bcd
mHb
b
0.1
mHb
Tor=0
(α)
0
Bcd
0.2
c
contribution
Bcd
Tor
mHb
0.05
y
0.95
d
probability
0
1
ỹ
y
4%
3%
2%
1%
0%
RMSE
(scaling error)
0.05
y
0.95
e
Null model
f
g
h
i
Scaling decoder
j
k
l

**Fig. S10-continued**

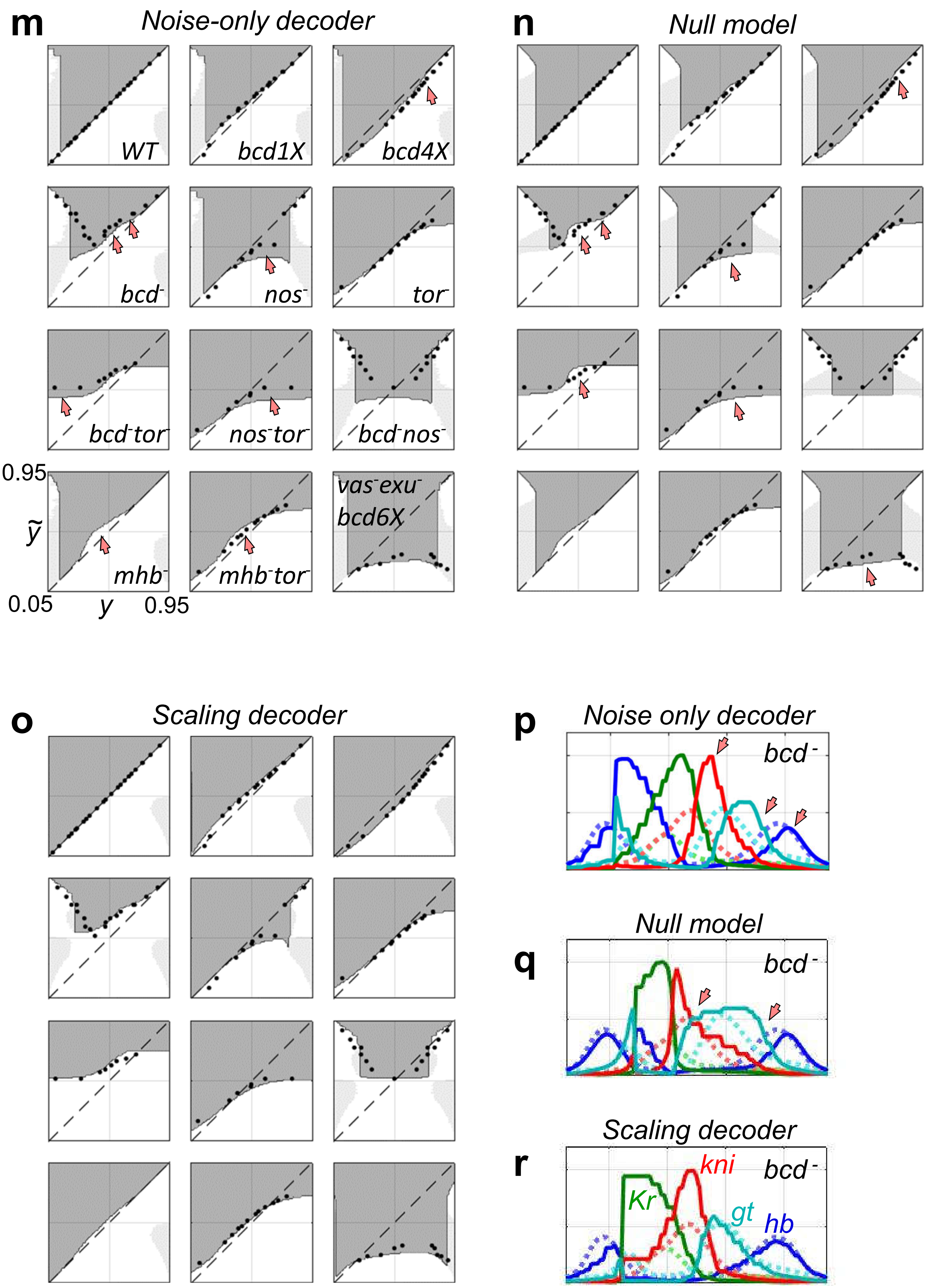

### Fig. S10-continued

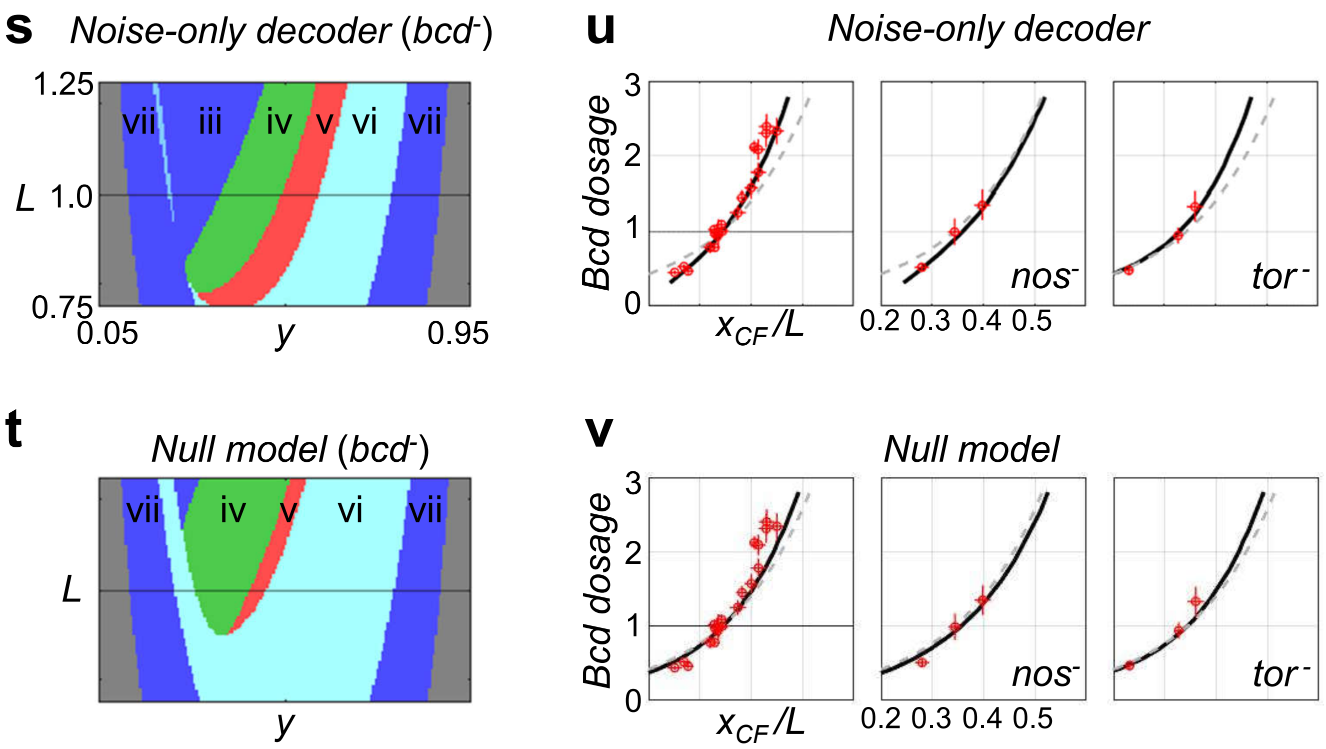

### Fig. S11

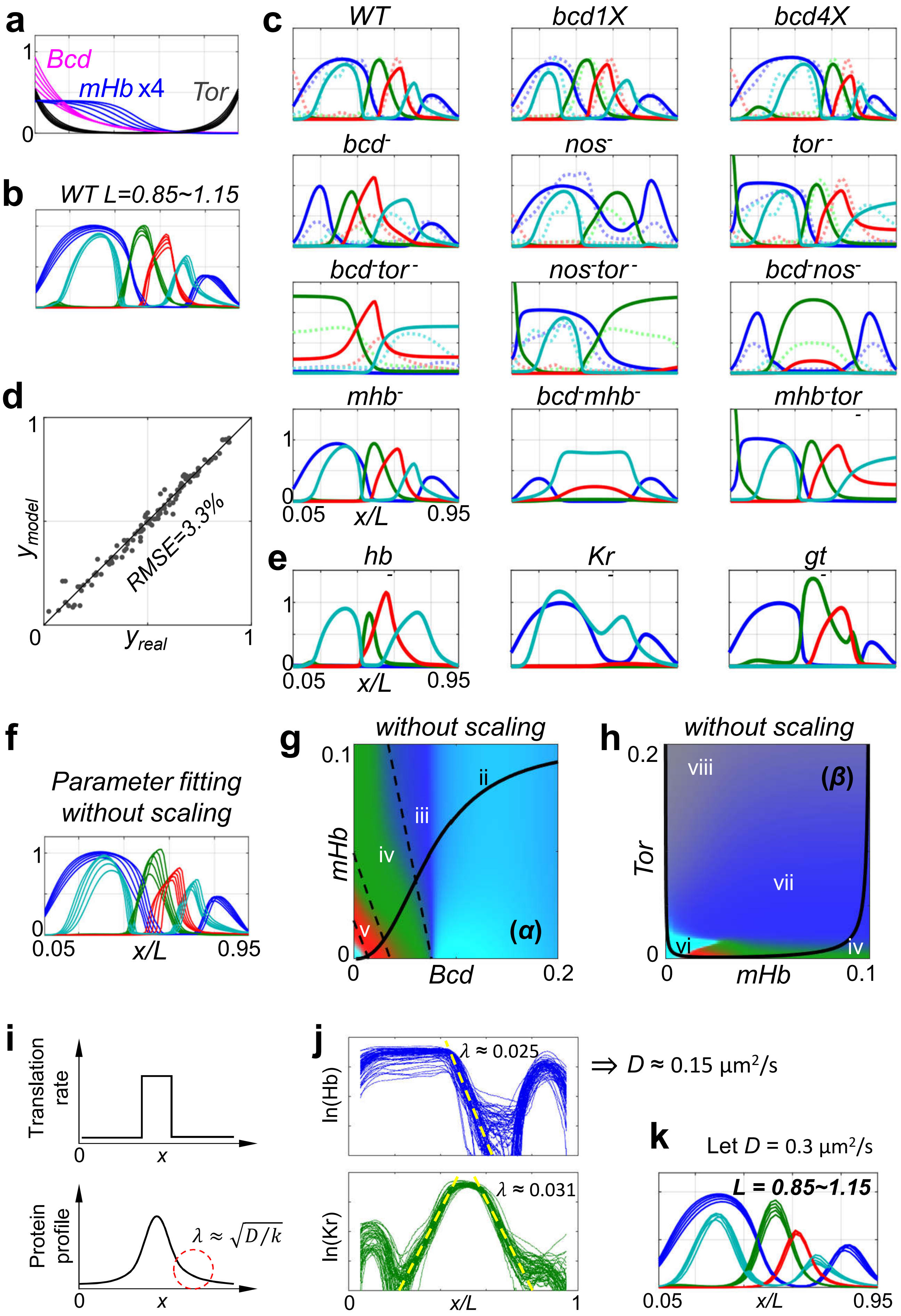

a
Bcd
mHb x4
Tor
b
WT L=0.85~1.15
c
WT
bcd1X
bcd4X
bcd-
nos-
tor-
bcd-tor-
nos-tor-
bcd-nos-
mhb-
bcd-mhb-
mhb-tor-
0.05
x/L
0.95
d
RMSE=3.3%
y_model
y_real
e
hb
Kr
gt
f
Parameter fitting
without scaling
g
without scaling
mHb
Bcd
(α)
h
without scaling
Tor
mHb
(β)
i
Translation rate
Protein profile
x
λ ≈ √(D/k)
j
ln(Hb)
ln(Kr)
λ ≈ 0.025
λ ≈ 0.031
⇒ D ≈ 0.15 μm²/s
x/L
k
Let D = 0.3 μm²/s
L = 0.85~1.15

#### Fig. S12

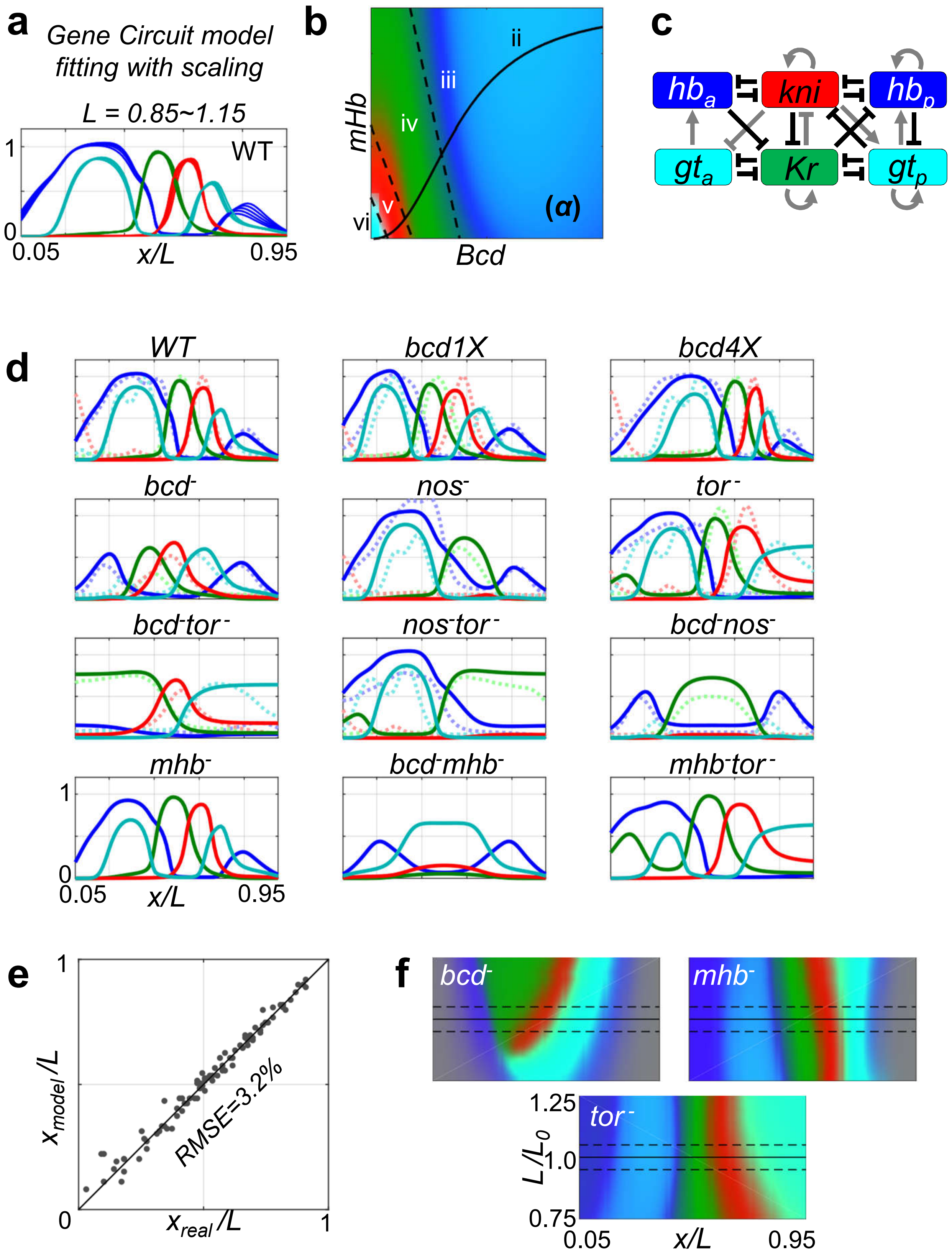

a
Gene Circuit model
fitting with scaling
L = 0.85~1.15
WT
1
0
0.05
x/L
0.95
b
mHb
Bcd
ii
iii
iv
v
vi
(α)
c
hb_a
kni
hb_p
gt_a
Kr
gt_p
d
WT
bcd1X
bcd4X
bcd-
nos-
tor-
bcd-tor-
nos-tor-
bcd-nos-
mhb-
bcd-mhb-
mhb-tor-
1
0
0.05
x/L
0.95
e
1
0
x_model/L
x_real/L
1
RMSE=3.2%
f
bcd-
mhb-
tor-
1.25
L/L_0
1.0
0.75
0.05
x/L
0.95

**Fig. S13**

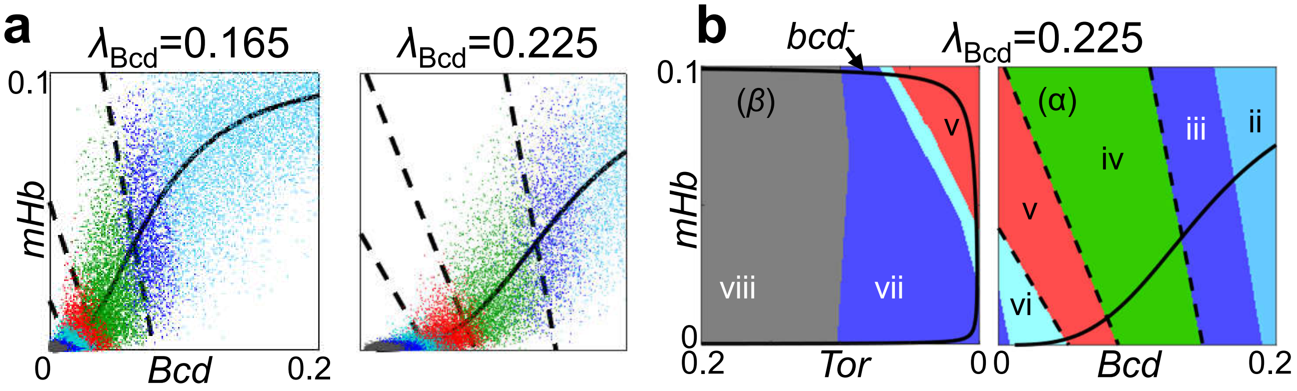

a
λBcd=0.165
λBcd=0.225
0.1
mHb
0
Bcd
0.2
b
bcd-
λBcd=0.225
0.1
(β)
v
viii
vii
0
0.2
Tor
0
(α)
iv
iii
ii
v
vi
0
Bcd
0.2
